# Landscape-scale navigation unlocks antibody CDR structural logic for AI-guided rescue and therapeutic optimization

**DOI:** 10.64898/2026.04.21.719857

**Authors:** Changju Chun, Byeong-Kwon Sohn, Heesoo Ki, Ji Hye Jo, Hyoung-Tae An, Jinseo Park, Jiyu Lee, Sooyeon Choi, Jayeon Choi, Hyeyoung Cho, Seong Beom Lee, Booyoung Yu, Chae Young Lee, Ji Eun Kim, Yu-jin Ban, Young-Yoon Choi, Byoungsan Choi, Hongwon Lee, Junho Chung, Minkyung Baek, Tae-Young Yoon

## Abstract

While AI offers transformative potential for therapeutic antibody design, the lack of ground-truth data fundamentally constrains our ability to model the epistatic topology of fitness landscapes. Here, we establish a high-throughput workflow to characterize tens of thousands of antibody variants per week with gold-standard biophysical precision. By combinatorially assembling functional variants from deep mutational scanning, we charted antibody fitness landscapes comprising over 17,000 data points, which revealed an extremely rugged, non-navigable epistatic topology. Yet, navigating at this unprecedented scale enabled the discovery of rare peak clusters exhibiting simultaneous enhancements in affinity and productivity. Strikingly, ProteinMPNN predicted the CDR-dependent productivity landscape with remarkable accuracy, suggesting that sequence-structure compatibility within CDRs gates cellular productivity. This insight enabled a structure-guided rescue strategy combining AlphaFold3 and ProteinMPNN, which successfully restored the cellular productivity of high-affinity, low-productivity clones via single amino acid substitutions. Two elite variants drawn directly from peak clusters further demonstrated 20- to 100-fold *in vivo* efficacy gains in a murine psoriasis model. Our findings establish CDR structural fitness as a fundamental determinant of antibody cellular productivity and validate landscape-scale navigation as a powerful framework for therapeutic antibody optimization.

## Introduction

Antibodies are the primary effectors of adaptive immunity, achieving exquisite specificity for diverse molecular targets through highly evolved recognition interfaces^1–4^. To engage a nearly infinite antigenic universe, antibodies utilize conformational plasticity within their complementarity-determining regions (CDRs)^5–7^, an evolutionary strategy that expands functional diversity without a commensurate increase in genomic encoding^2,8,9^. The theoretical sequence space of the CDRs is astronomically large, and identifying optimal therapeutic binders thus requires navigating a vast and largely uncharted fitness landscape.

The recent emergence of antibody design via artificial intelligence (AI) offers the potential to access regions of sequence space that natural evolution has not yet reached^10–15^. However, the current reality of *de novo* design is hindered by limited success rates, largely attributed to the inability of models to accurately predict CDR conformations and their interaction with antigens^13,16,17^. Beyond binding affinity, therapeutic success hinges on the parallel optimization of developability traits, including cellular productivity and thermal stability^18–20^. A recurring bottleneck in AI-driven design is the dramatic loss of expression in mammalian cells, which frequently nullifies high-affinity candidates^21,22^. This expression failure is difficult to reconcile with traditional view of CDRs as flexible loops whose conformations are primarily shaped by antigen engagement^23^, a view that largely precludes the existence of intrinsic structural constraints on antibody productivity. Consequently, a fundamental prerequisite for advancing AI-driven antibody design would be the generation of ground-truth, high-resolution learning materials that map the intertwined relationship between affinity and developability across broad sequence diversity^24–26^. Charting these multi-dimensional fitness landscapes would reveal the epistatic topology of sequence variation, potentially answering whether affinity and developability represent an inherent trade-off or if they permit simultaneous optimization through strategic navigation of sequence space.

To date, few attempts have been made to delineate these landscapes, and their underlying topology remains largely unexplored^27–30^. The lack of high-fidelity maps represents a long-standing technological impasse, which is the inability to combine the throughput of library-scale screening with the quantitative rigor of classical biophysics. Conventional immunization and library-display technologies emphasize scale but typically assess binders through low-resolution, binary classifications^31,32^. Increasing data quantitativeness in these systems requires complex calibration or modeling to account for cellular heterogeneity, which in turn significantly restricts the scope of sequence navigation^33–35^. At the other extreme, gold-standard biophysical methods, such as surface plasmon resonance (SPR) and differential scanning fluorometry (DSF), provide high precision but are bottlenecked by the requirement for moderate-scale expression and rigorous purification, severely limiting throughput^36–39^.

To bridge this technological gap, we developed an alternative experimental workflow that characterizes antibody landscapes at the scale of tens of thousands of clones while maintaining the quantitative precision of classical biophysics. We leveraged the Single-molecule Protein Interaction Detection (SPID) platform, inspired by single-molecule pull-down techniques and previously used for sensitive detection of protein complexes in clinical specimens^40–43^. This adaptation enables the direct determination of dissociation constants (*K*_D_) using single-molecule imaging and counting without requiring the purification of either antibody or antigen, thereby significantly reducing experimental burden^44^. As a result, we generated high-resolution, quantitative data for tens of thousands of clones per week, while maintaining a level of precision on par with the standard biophysics methods^36,37^. Crucially, this workflow simultaneously quantifies cellular productivity and thermal stability, providing a multi-dimensional view of each variant’s fitness. Using this workflow, we performed deep mutational scanning (DMS) to map the effects of all possible single-residue substitutions across the CDRs of ten therapeutic antibodies^45,46^. This comprehensive dataset revealed that buried CDR residues significantly influence antibody cellular productivity in a depth-dependent manner, reminiscent of folded protein domains^33,35,47^. To explore the sequence space beyond single-residue variation, we demarcated functional variations from destructive ones based on this high-resolution DMS data. The combinatorial assembly of these variants for the cytokine-neutralizing antibody adalimumab revealed a highly rugged fitness landscape dominated by negative epistasis. However, our mapping scale allowed us to identify rare clusters of positive epistasis that simultaneously enhanced both affinity and productivity.

Strikingly, scoring these sequences with the deep-learning model ProteinMPNN^48^ showed unexpectedly high precision in predicting the productivity landscapes, suggesting that sequence-structure compatibility within CDRs is a core determinant of antibody productivity in mammalian cells. Leveraging this insight, we integrated structure-prediction engines like AlphaFold3^49^ with ProteinMPNN to redesign the CDR sequences of high-affinity, low-productivity clones, successfully restoring productivity without compromising binding. Finally, the administration of two variants isolated from the positive-epistasis cluster achieved a 20- to 100-fold dose reduction in a psoriasis-like mouse model, with this enhanced potency attributable to an extended antibody–antigen complex lifetime rather than improved equilibrium affinity. Collectively, these findings reveal the intrinsic structural logic of CDRs underlying antibody cellular productivity and demonstrate that landscape-scale navigation can deliver therapeutic antibodies beyond the reach of conventional, incremental optimization.

## Results

### Generating precision antibody data at scale with SPID

We tailored the SPID platform to quantify antibody–antigen interactions and developability properties (cellular productivity and thermal stability) at scale, without compromising data precision. For validation, we focused on two clinically established therapeutic antibodies, trastuzumab and adalimumab, which target human epidermal growth factor receptor 2 (HER2) and tumor necrosis factor-α (TNF-α), respectively^50–52^. We evaluated multiple antibody formats—single-chain variable fragment (scFv), single-chain fragment antigen-binding (scFab) and full-length immunoglobulin G (IgG)—each fused at the C-terminus to a red fluorescent protein (RFP, specifically mScarlet). Our findings confirmed that the SPID workflow is fully compatible with all of these formats (see Methods for vector designs) (Fig. 1a, left). To enable CDR sequence editing, we introduced silent mutations that created type II restriction enzyme sites flanking the target CDRs, together with multiple tandem stop codons to prevent antibody production from undigested vectors (Fig. 1a and Extended Data Fig. 1a). Double-stranded DNA inserts encoding CDRs and linkers were assembled and ligated into the digested backbone (Extended Data Fig. 1b-d). As little as ∼100 ng of the resulting vectors was sufficient to drive expression and secretion of antibodies in mammalian cells, yielding nanomolar antibody concentrations in the culture media and eliminating the need for vector amplification (Extended Data Fig. 1e,f).

**Fig. 1.**
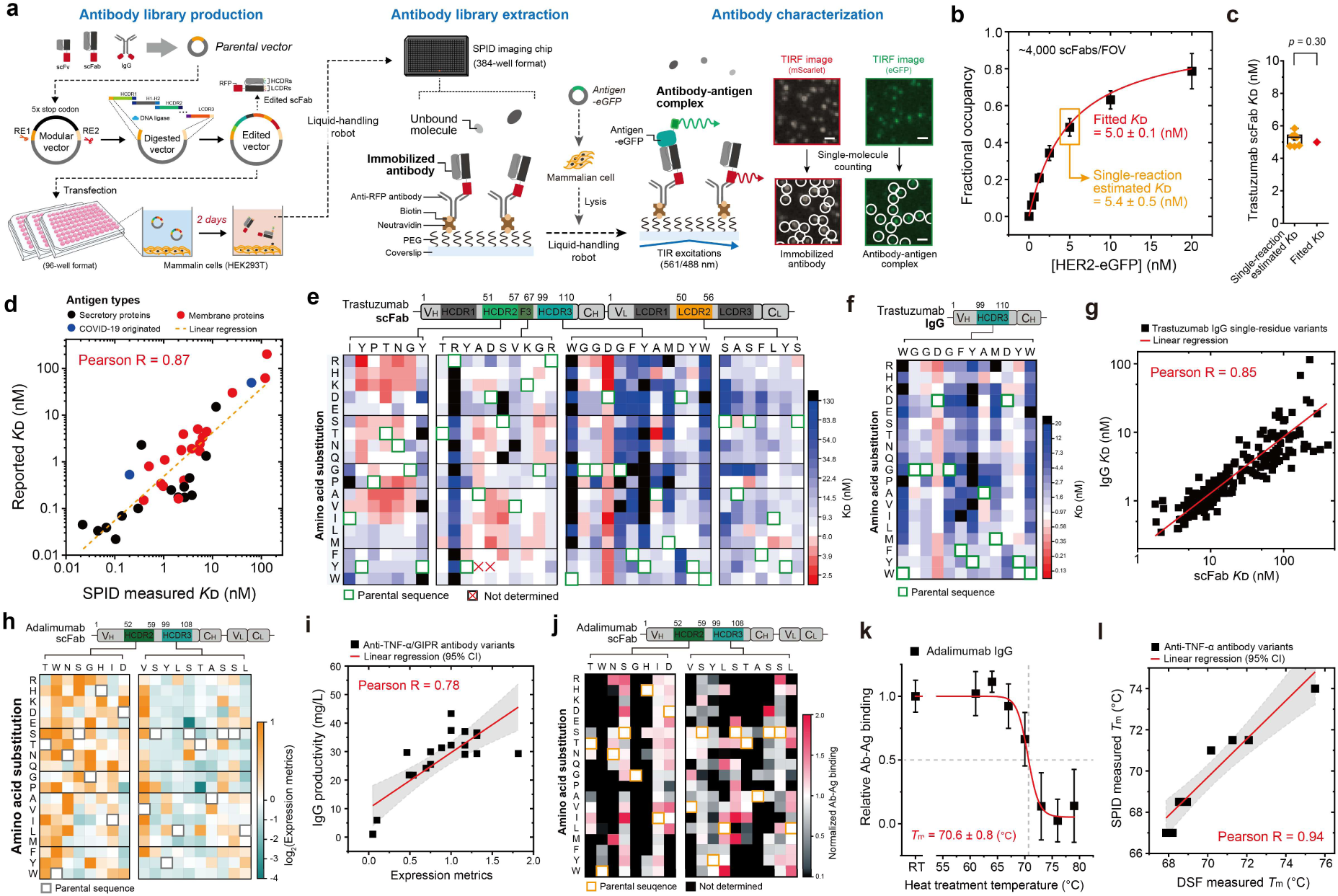
The SPID platform allows high-throughput, multi-parametric profiling of antibody-antigen interaction landscapes. **a**, Schematic overview of SPID-based rapid antibody library production and antibody characterization. **b**, Binding curves for HER2-eGFP to trastuzumab scFab used for *K*_D_ fitting. **c**, Comparison of *K_D_* values for the trastuzumab scFab-HER2 interaction obtained by single-reaction estimates and curve fitting (two-sided one-sample t-test, *p*=0.30). **d**, Correlation of *K*_D_ values measured using SPID and previously reported values of original antibodies. A total of 36 distinct antibody sequence-based scFabs were analyzed to assess the correlation between their measured affinities and the reported IgG values. **e**, Deep mutational scanning (DMS) of *K*_D_ changes in trastuzumab scFab-HER2 interactions from single-residue variations in CDRs (HCDR2, HCDR3, and LCDR2) and the framework region 3 (F3). **f**, Affinity DMS heatmap of trastuzumab IgG HCDR3 variants corresponding to (**e**). **g,** Comparison of *K*_D_ values measured via SPID between trastuzumab scFab and IgG single-residue variants. **h**, DMS of expression metrics for adalimumab scFab single-residue variants. **i**, Correlation between SPID-measured expression metrics and productivity from large-scale (up to 10 liters) cultures of anti-TNF-α or anti-GIPR antibodies in HEK293F cells. **j**, DMS of adalimumab scFab-TNF-α binding after 60 °C heat treatment (Ab: antibody, Ag: antigen). **k,** Changes of TNF-α binding for adalimumab IgG as a function of heat treatment temperature (*T*ₘ: Melting temperature). **l,** Correlation between *T*ₘ values determined by SPID and the Differential Scanning Fluorimetry (DSF) method.

Using a liquid-handling robot, we directly transferred conditioned media into individual wells of the imaging chip, patterned to match a 384-well plate format (Fig. 1a, middle). Secreted antibodies in the media were then directly pulled down onto a polyethylene glycol (PEG)-coated surface via anti-RFP antibodies, while virtually eliminating non-specific adsorption of other proteins in the media (Extended Data Fig. 1g). This allowed us to bypass conventional purification steps^32^. The number of immobilized antibodies remained stable even at higher cell extract or media concentrations, indicating minimal interference from the cellular proteome (Extended Data Fig. 1h)^41,43^.

To present antigens in a near-native context, we fused HER2 or TNF-α to enhanced green fluorescent protein (eGFP) at their C-termini and expressed these constructs in mammalian cells (Fig. 1a, right). Guided by our prior studies, we employed a lysis protocol with non-ionic or cholesterol-mimicking detergents that preserved native conformations and tyrosine kinase activities of growth factor receptor complexes^41,42^. Crude lysates were diluted and introduced to reconstitute antibody–antigen interactions (Fig. 1a, right and Extended Data Fig. 1i-k).

Antibody-antigen complexes appeared as stable, diffraction-limited spots under single-molecule total internal reflection (TIR) fluorescence imaging (Fig. 1a, right insets). Control experiments confirmed both the specificity of antibody–antigen interactions within dense lysate environments and the maintenance of constant fractional occupancy when multiple CDRs were simultaneously substituted (Extended Data Fig. 1l,m).

When we varied antigen concentration at a fixed antibody immobilization level, the resulting fractional occupancy values closely followed the Hill equation, yielding dissociation constants (*K*_D_) (Fig. 1b and Extended Data Fig. 2a). Across independent biological replicates (60 total measurements), coefficients of variation were below 5%, highlighting the robustness and reproducibility of the SPID workflow (Extended Data Fig. 2b). We further tested whether reliable *K*_D_ estimates could be obtained from single antigen concentrations, bypassing full titrations. Individual single-point measurements produced a narrow *K*_D_ distributions (Fig. 1b,c), demonstrating that SPID can deliver precise affinity estimates from single-condition assays (Extended Data Fig. 2c-g)^53–56^.

To assess the broader applicability of this approach, we cloned 36 additional therapeutic antibodies into scFab or IgG vectors and profiled their antigen interactions using the SPID workflow. Across this panel, *K*_D_ values measured by SPID showed strong correlation with those determined by SPR or biolayer interferometry (Fig. 1d)^51,53–90^. Notably, most antigens were assayed in their full-length, unpurified forms, with intact transmembrane domains in the case of membrane proteins (Supplementary Table 1), maintaining the structural context of native epitopes (Extended Data Fig. 2h-j)^58^.

To demonstrate that SPID can support DMS-scale library screening, we performed pilot DMS on trastuzumab and adalimumab, systematically substituting each CDR position with all naturally occurring amino acids, excluding cysteine (Fig. 1e-l and Fig. 2a). A predominance of dark-blue and black positions in heavy chain CDR3 (HCDR3) underscored its critical role in antigen recognition for both antibodies, whereas white points interspersed with modest red and light-blue shifts indicated largely incremental affinity changes at many other sites (Fig. 1e and Fig. 2a)^91^. When DMS was performed for HCDR3 of an IgG construct of trastuzumab, despite differences in absolute *K*_D_ values due to avidity effects of bivalent binding (5.0 nM versus 0.8 nM), *K*_D_ trends for IgG variants correlated well with those of the corresponding scFab variants (Fig. 1e-g), reconfirming that SPID affords consistent affinity measurements across antibody formats.

**Fig. 2.**
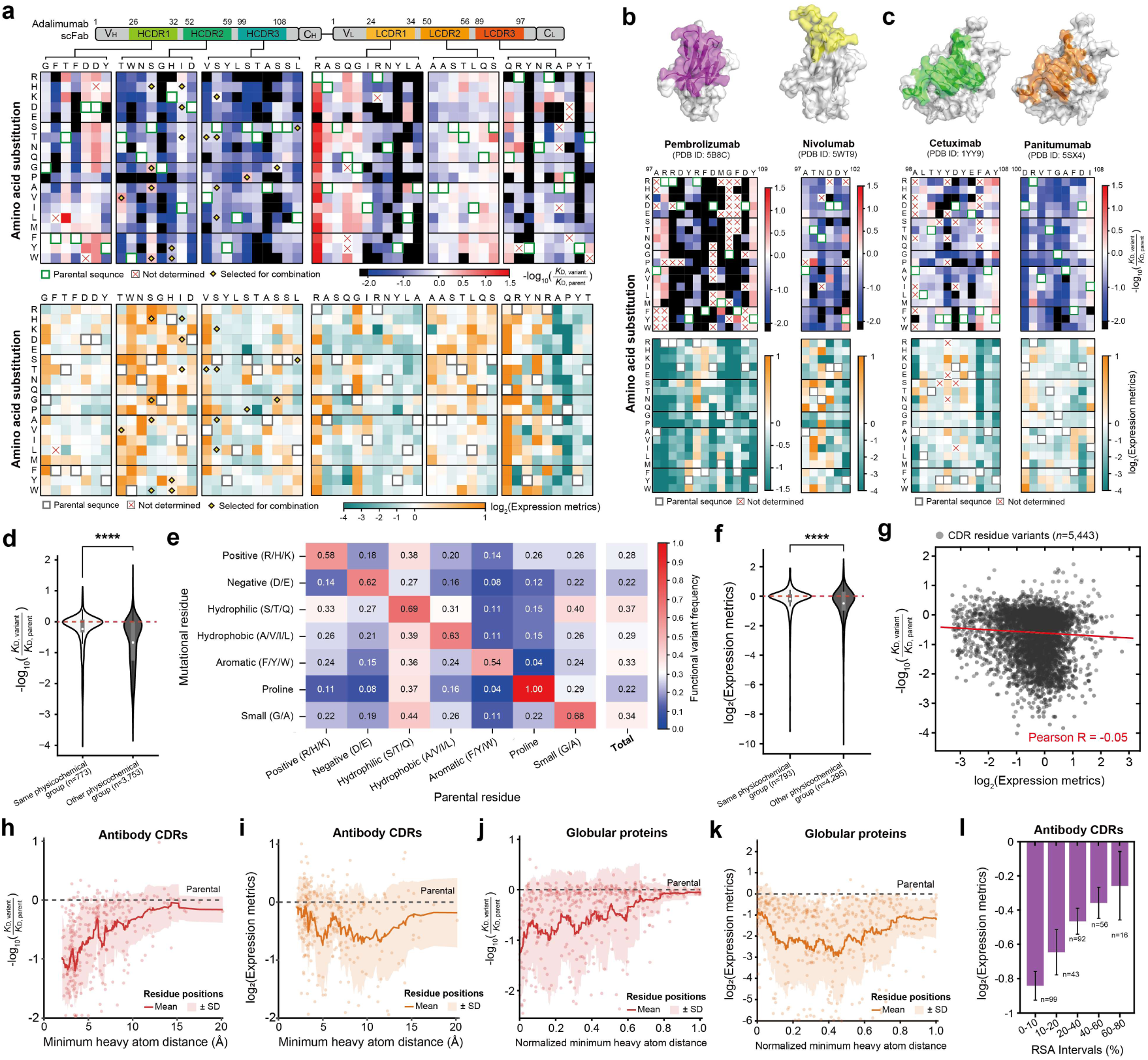
Comprehensive deep mutational scanning of therapeutic antibodies reveals the biophysical logic of CDR function. **a**, Comprehensive DMS heatmaps for the six CDRs of adalimumab scFab (upper: relative fold changes in *K*_D_; lower: relative fold changes in expression metrics). **b,c,** Comparison of DMS profiles between antibody pairs targeting the same antigen. Structures of (**b**) PD-1 and (**c**) EGFR are shown with their respective epitopes highlighted in color. **d-f**, Impact of physicochemical substitutions on functional properties across 10 FDA-approved therapeutic antibodies (see **Extended Data** Fig. 4 for the full list). (**d**) Distribution of affinity changes for substitutions within the same or different physicochemical groups (two-sided Mann-Whitney U-test), (**e**) Functional variant frequency categorized by the physicochemical property of the substituted residue, (**f**) Distribution of expression changes (two-sided Mann-Whitney U-test). (**d** and **f**) The dashed line indicates the threshold for functional variants. **g,** Correlation between affinity and expression metrics of antibody variants (*n*=5,443). **h-k,** Mean functional changes as a function of the minimum heavy atom distance from the binding interface. (**h**) Affinity and (**i**) expression metrics averaged across 324 CDR residues from 10 therapeutic antibodies. (**j**) Affinity and (**k**) expression metrics averaged across globular proteins (GRB2, KRAS, PSD95, SARS-CoV-2 RBD). **l,** Mean fold change in expression metrics for 324 CDR residues categorized by relative solvent accessibility (RSA).

Distinct CDR sequences can also markedly influence antibody cellular productivity (Fig. 1h). To capture this complexity, we first confirmed that RFP counts scaled proportionally with the amount of antibody introduced (Extended Data Fig. 1h and Extended Data Fig. 3a), and that these expression metrics were reproducible across biological replicates, independent of antigen affinity (Extended Data Fig. 3b-d). We compared SPID-derived expression metrics for 19 IgG antibodies (anti-TNF-α and anti-glucose-dependent insulinotropic polypeptide (GIP) receptor antibodies) with production yields obtained from larger-scale HEK293F cultures ranging from 40 mL to 10 L (Fig. 1i)^92^. The resulting high correlation confirmed that the expression metrics assessed by SPID can provide early, small-scale indicators of cellular productivity.

We next attempted to assess thermal stability within the SPID workflow. Antibody-containing media were exposed to a 10-min heat pulse at defined temperatures, after which antigen binding was measured (Extended Data Fig. 3e). The ratio of antigen-binding occupancy with and without heat treatment (for example, at 60 °C) served as a robust measure of relative thermal stability for each clone (Fig. 1j). Moreover, by titrating the heat-treatment temperature, we were able to determine melting temperatures (*T*_m_) (Fig. 1k), which showed strong correlation with those obtained by conventional DSF following large-scale antibody expression and purification (Fig. 1l)^93^. Collectively, these results demonstrate the extension of the SPID platform into a multi-parametric analytical tool. By enabling the simultaneous assessment of antigen affinity and cellular productivity via two-color single-molecule imaging within a single well dedicated to each clone, the platform seamlessly facilitates the profiling of landscape-scale libraries encompassing tens of thousands of variants. Additionally, *T*_m_ determination—which requires sampling multiple temperature points per clone—was subsequently applied as a rigorous quality filter for prioritized variants within our workflow.

### Deep mutational scanning of therapeutic antibody CDRs

Employing the high-throughput capabilities of the SPID workflow, we conducted DMS for ten FDA-approved therapeutic antibodies with known antigen-binding structures (Fig. 2 and Extended Data Fig. 4)^54–66^. This effort generated a comprehensive dataset encompassing 5,832 substitutions across 324 residues within 33 CDRs (excluding cysteine variations) (Supplementary Dataset 1). While the SPID platform provided absolute *K*_D_ values ranging from the sub-nanomolar to nanomolar scale, we visualized the data as relative fold changes in *K*_D_ to facilitate the identification of emergent patterns (Fig. 2a-c). Correspondingly, cellular expression metrics were normalized to those of the parental sequences. Despite shared characteristics, such as the prevalence of affinity-abolishing mutations (indicated in black) within the HCDR3, each antibody exhibited a distinct DMS topography (Fig. 2a-c and Extended Data Fig. 4a). For instance, although pembrolizumab and nivolumab target overlapping epitopes on programmed cell death protein 1 (PD-1)^63,94^, they possess different HCDR3 lengths and display divergent DMS profiles (Fig. 2b). Notably, a significant proportion of single-residue substitutions in the pembrolizumab HCDR3 resulted in markedly diminished expression levels and rendered *K*_D_ determination infeasible (marked with ‘x’). Similarly, cetuximab and panitumumab, which target overlapping EGFR epitopes^65,95^, showed distinct HCDR3 DMS maps and multiple expression-reducing variations (Fig. 2c). Collectively, these DMS data highlight the diverse fingerprints of CDRs that reflect the unique biophysical signatures of individual antibodies.

The majority of single-residue variations were found to impair rather than enhance antigen affinity. Consequently, we focused on affinity-maintaining or affinity-enhancing mutations (hereafter referred to as functional variants) and investigated their physicochemical properties (Supplementary Dataset 2). As anticipated, substitutions within the same physicochemical group (e.g., Lys to Arg or His) showed the highest probability of yielding functional variants (Fig. 2d). Notably, the frequency of these functional variants was higher for substitutions to hydrophilic residues (0.37) than for those to charged (Positive: 0.28, Negative: 0.22), hydrophobic (0.29), or aromatic residues (0.32) (Fig. 2e). In detail, serine and threonine exhibited the highest success rates of 0.42 and 0.37 respectively, for maintaining or enhancing affinity (Extended Data Fig. 5a). This suggests that within the sophisticated local context of the paratope, bulky or long-range residues, often considered nuclei for protein-protein interfaces, may not always be prioritized over smaller residues with hydrogen-bonding capabilities like serine and threonine for affinity enhancement.

Consistent with the trends observed for antigen affinity, substitutions within the same physicochemical group exhibited a higher degree of tolerance regarding cellular productivity (Fig. 2f, and Extended Data Fig 5b,c). Nevertheless, many single-residue variations led to substantial alterations in cellular productivity as observed for pembrolizumab and cetuximab (Fig. 2b,c). These observations align with prior reports in antibody engineering and AI-based design, demonstrating that even minor sequence variations within the CDRs can markedly influence antibody cellular productivity^33,35,47^. When fold changes in productivity were plotted against changes in antigen affinity, we observed no significant correlation, demonstrating that the principles governing antibody expression in mammalian cells are essentially orthogonal to those driving antigen recognition (Fig. 2g). Of note, this orthogonality could be of fundamental importance, as it implies that the fitness cliffs of productivity and affinity do not necessarily overlap, providing a window for the optimization of one property without an inherent trade-off in the other.

Finally, we investigated how general spatial patterns in CDR residues govern their respective contributions to affinity and cellular productivity. We classified the 324 residues based on their distance from the antigen, defined by the minimum heavy-atom distance in the antigen-bound structure, and calculated the functional changes for each residue (Supplementary Dataset 3). The mean loss in affinity was most significant for residues in the immediate proximity of the antigen and decayed with increasing distance (Fig. 2h). Conversely, the impact on productivity was minimal for these surface-exposed residues but exhibited a broad valley for CDR residues located between 5 and 10 Å from the interface (Fig. 2i). This observation was unexpected, as such patterns are typical of globular proteins, where buried residues are under stronger structural constraint than surface-exposed ones, a trend we reconfirmed by applying the same analysis to globular protein DMS data (Fig. 2j,k and Extended Data Fig. 5d-g)^33,35,47^. To further validate this, we reclassified the 324 residues based on relative solvent accessibility (RSA; defined by the ratio of the observed solvent-accessible surface area of a residue to its maximum possible value) rather than simple distance from the antigen (Supplementary Dataset 3)^96–98^. When the fold change in productivity was averaged across different RSA intervals, it indeed increased monotonically as RSA values declined (Fig. 2l and Extended Data Fig. 5h). Collectively, these results suggest that CDR residues contribute to mammalian cell expression in a depth-dependent manner characteristic of folded proteins, pointing to a defining role for the structural context of CDR sequences in cellular productivity, a hypothesis we systematically investigate below.

### Exploring fitness landscapes for adalimumab

Leveraging the large-scale, multi-parametric profiling capability established above, we next sought to chart antibody fitness landscapes beyond DMS, which primarily interrogates single-residue variation. To ensure the SPID platform reflected the therapeutically relevant binding mode, we confirmed that adalimumab preferentially binds to the biologically relevant trimeric TNF-α over the monomeric form, as evidenced by a rightward shift in the fluorescence distribution of single adalimumab-TNF-α complexes (Extended Data Fig. 6a). Building on the multi-parametric DMS maps for adalimumab HCDR2 and HCDR3, we selected functional substitutions as those that maintained or improved both antigen affinity and productivity relative to the parental adalimumab. These functional substitutions were distributed across nine positions spanning HCDR2 and HCDR3 (Fig. 3a). Importantly, this filtering step dramatically reduced the effective search space, as the subsequent systematic recombination of these substitutions yielded a target library of 9,600 variant clones. Using a single SPID imaging system over a four-week period, we successfully characterized both binding affinity and cellular productivity for 9,517 clones (Supplementary Dataset 4), marking a 99.1% success rate for the workflow.

**Fig. 3.**
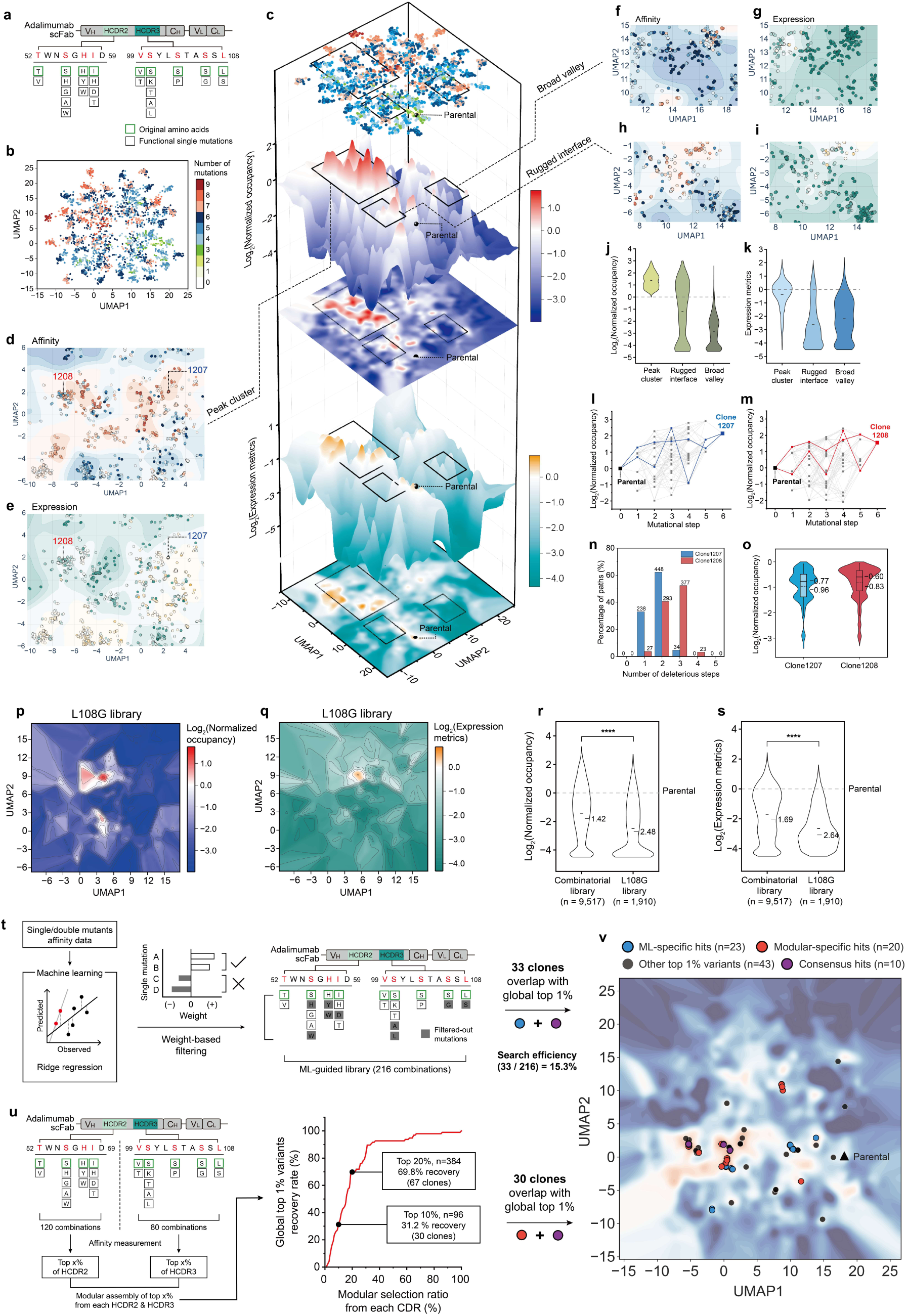
Topographical fitness landscapes of the combinatorial antibody library and efficient strategies for identifying global optimum variants. **a,** Functional single-amino-acid substitutions were introduced into the HCDR2 and HCDR3 loops of adalimumab based on affinity deep mutational scanning (DMS) profiling to construct a combinatorial sequence library. **b,** Two-dimensional (2D) UMAP representation of ∼9,600 variants, vectorized by mutation type, position, and physicochemical descriptors. Each dot denotes a variant colored by mutation count. **c,** Three-dimensional fitness landscapes projected on the UMAP x–y plane, with SPID-derived metrics on the z-axis. The upper surface (red–white–blue) indicates affinity (relative occupancy), and the lower surface (orange–white–teal) shows expression metrics, both plotted on a log₂-transformed z-axis. **d–i,** Representative magnified regions highlighting three characteristic topographies: (**d** and **e**) peak cluster, (**f** and **g**) broad valley, and (**h** and **i**) rugged interface. Characteristic regions were defined by combining K-nearest neighbors (KNN, k=30) and density-based spatial clustering (DBSCAN). The peak cluster comprises densely grouped high-affinity variants, the broad valley contains contiguous low-affinity basins, and the rugged interface represents areas of high local variability in affinity. Detailed classification criteria are provided in the Methods section. **j,k,** Violin plots comparing (**j**) relative occupancy and (**k**) expression metrics across these topographical classes. **l,m,** Evolutionary trajectories from the parental sequence to optimized clones (**l**) 1207 and (**m**) 1208 within the peak cluster, with two representative pathways highlighted. **n,** Distribution of all possible evolutionary pathways for clone1207 (blue) and 1208 (red) by the number of deleterious steps. **o,** Violin plot illustrating the magnitude of stepwise affinity loss during deleterious steps across the pathways for both clones. **p,q,** 2D contour plots showing (**p**) normalized occupancy and (**q**) expression of variants containing the L108G mutation, colored as in (**c**). **r,s,** Violin plots comparing (**r**) normalized occupancy and (**s**) expression between the full and L108G-restricted libraries. **t,** Machine learning (ML)-guided library design based on additive mutation weights from ridge regression of single/double mutants. Filtering recovered 216 combinations, including 33 clones overlapping with the global top 1%. **u,** Schematic representation and quantitative evaluation of the modular assembly strategy. The line graph illustrates the relationship between the modular selection ratio and the recovery rate of global top 1% variants. **v,** Projection of global top 1% variants on the UMAP affinity landscape, color-coded as ML-specific (blue), modular-specific (red), consensus (purple), and other top 1% variants (gray).

To visualize the resulting fitness landscapes, we embedded each clone as a vector encoding the physicochemical properties of the substituted residues across the nine targeted sites (Extended Data Fig. 6b). These vectors were projected onto a two-dimensional plane using density-preserving Uniform Manifold Approximation and Projection (densMAP). In the resulting embedding, clones with higher mutation counts diverged progressively from the parental sequence, and the average distance between clones differing by a single mutation increased systematically with additional mutations, confirming that the projection preserves the topology of the sequence space (Fig. 3b and Extended Data Fig. 6c,d). By overlaying antigen affinity and productivity metrics onto this embedding, we resolved the fitness landscapes across the 9,517-clone sequence space (Fig. 3c). Because all variants target the same antigen and epitope, we used antigen occupancy normalized to the parental clone (rather than *K*_D_ values) as the affinity measure, minimizing error propagation from Hill equation fitting. Notably, despite restricting our library to functional building blocks—substitutions that were neutral or beneficial in the parental DMS maps—both the affinity and productivity landscapes were characterized by pervasive valleys. Specifically, 73.5% and 81.7% of the 9,517 combinations exhibited a loss of affinity or productivity, respectively, compared to the parental clone. This global topography reveals dominant negative epistasis among mutations that are individually functional.

Within the high-dimensional sequence space, we identified regions characterized by clustered local peaks in affinity and productivity (Fig. 3d-i and Extended Data Fig. 6e-j). A magnified analysis of one of these peak clusters revealed that affinity and productivity peaks, despite occasional overlaps, are generally disparate, reinforcing the biophysical decoupling of these traits previously observed in the DMS data (Fig. 3d,e and Extended Data Fig. 6l). Closer investigation into regions of pervasive negative epistasis (broad valleys) and sharp transitions between fitness states (rugged interfaces) further reconfirmed the persistent discordance between the affinity and productivity landscapes (Fig. 3f-i). These discrete peaks were often directly adjacent to clones with nearly abolished fitness, attesting to the highly rugged architectures of both landscapes. Mutational profiling indicated that peak clusters and broad valleys are enriched with distinct sets of mutations, while the interfacial regions exhibit intermediate profiles (Fig. 3j,k and Extended Data Fig. 6e-k).

We selected two clones from the main peak cluster marked in Figure 3d, referred to as 1207 and 1208, each harboring six substitutions while maintaining thermal stability. Given these six substitutions, there are 720 (6!) possible mutational trajectories leading from the parental sequence to either variant (Fig. 3l,m). Strikingly, pathway analysis revealed that every possible trajectory to 1207 and 1208 traversed at least one intermediate step characterized by a decrease in affinity, with the median value indicating a 50% loss in antigen occupancy (Fig. 3n,o). Thus, even within a library of individually functional substitutions, the landscape does not permit a strictly monotonic route to these high-fitness variants. This inherent non-navigability implies that incremental optimization strategies are unlikely to reach clones like 1207 or 1208, underscores the necessity of mapping fitness landscapes at scale with quantitative precision.

We further evaluated the composition and navigability of our search space to determine whether our DMS-guided combinatorial screening could be made more efficient in future applications. First, to test whether the DMS-based pre-filtering was necessary, we constructed a parallel library of 1,920 combinations in which all variants carried the deleterious L108G mutation alongside a subset of functional variants at the remaining positions (Fig. 3p,q, and Extended Data Fig. 7a-c). While a few fitness peaks remained, the probability of negative epistatic interactions between building blocks increased significantly, resulting in a marked expansion of void valley regions. Consequently, the probability of achieving positive epistasis was dramatically reduced upon the inclusion of a non-functional variant, supporting our strategy of recombining only individually functional single-residue substitutions (Fig. 3r,s).

We then explored strategies to navigate this functional space more efficiently. Previous studies have utilized machine learning to predict epistatic effects from low-order mutational combinations^99^, and our library already contained all single- and double-mutant combinations within the nine targeted positions. By applying ridge regression training to this single- and double-mutant dataset, we refined our amino-acid selection at each position beyond the initial DMS criteria (Extended Data Fig. 7d). Evaluating a total of 216 combinations within this refined subset recovered 33 of the top 1% of clones from the global library (which comprised 96 clones in total and exhibited five- to eight-fold increases in the antigen occupancy; Supplementary Dataset 5), representing a 15-fold improvement in hit rate over unbiased sampling from the pre-filtered library (Fig. 3t). We next designed a modular, divide-and-conquer approach, in which combinations within individual CDRs were first evaluated (120 and 80 combinations for HCDR2 and HCDR3, respectively) and the top-performing intra-CDR combinations were subsequently assembled (Fig. 3u). When assessing the recovery rate of the global top 1% clones while varying the selection threshold, we observed a ∼30% recovery when crossing the top 10% of combinations for HCDR2 and HCDR3. Expanding the threshold to the top 20% (yielding a total of 384 combinations) markedly increased the recovery rate to 70% (Fig. 3u and Extended Data Fig. 7e,f). Taken together, both ridge-regression refinement and modular assembly preserve landscape-informed decision making while substantially reducing screening burden, providing practical paths from library-scale data to high-fitness variants (Fig. 3v).

### AI-driven prediction and rescue of antibody cellular productivity

Prompted by our observation that buried CDR residues influence cellular expression similarly to those in globular proteins (Fig. 2h-k), we sought to extend this logic to our mapped antibody fitness landscapes. ProteinMPNN, a deep learning model originally developed for structure-based sequence design^48^, inherently learns the compatibility between amino acid sequences and their backbone geometry. We therefore repurposed it as a scoring tool, computing a Compatibility Score for each variant as the mean negative log-probability of variant’s CDR sequence given the adalimumab backbone structure (PDB ID: 3WD5, antigen excluded), normalized relative to wild-type such that positive values indicate improved structural compatibility.

Strikingly, normalized Compatibility Scores correlated consistently with antibody productivity levels across the full 9,517-variant library (Fig. 4a-c). This finding suggests that sequence-structure compatibility within CDRs is a core determinant of antibody cellular productivity. Examination of the bottom 10% of productivity revealed that substitutions at T52, S55, H57, and I58 in HCDR2, and S100, S103, S106, and L108 in HCDR3 were particularly deleterious (Fig. 4d and Extended Data Fig. 8d). Structural analysis reveals that these residues form a dense network of interactions with both the antibody framework and adjacent CDR loops (Fig. 4e-h, Extended Data Fig. 8e-h, and Supplementary Note 1), consistent with the interpretation that CDR sequence must be compatible with the surrounding structural context to support high productivity. In contrast, Compatibility Scores computed on the full antibody-antigen complex showed only marginal correlation with the affinity landscape (Extended Data Fig. 8a-c), indicating that ProteinMPNN does not predict affinity as effectively as it predicts productivity.

**Fig. 4.**
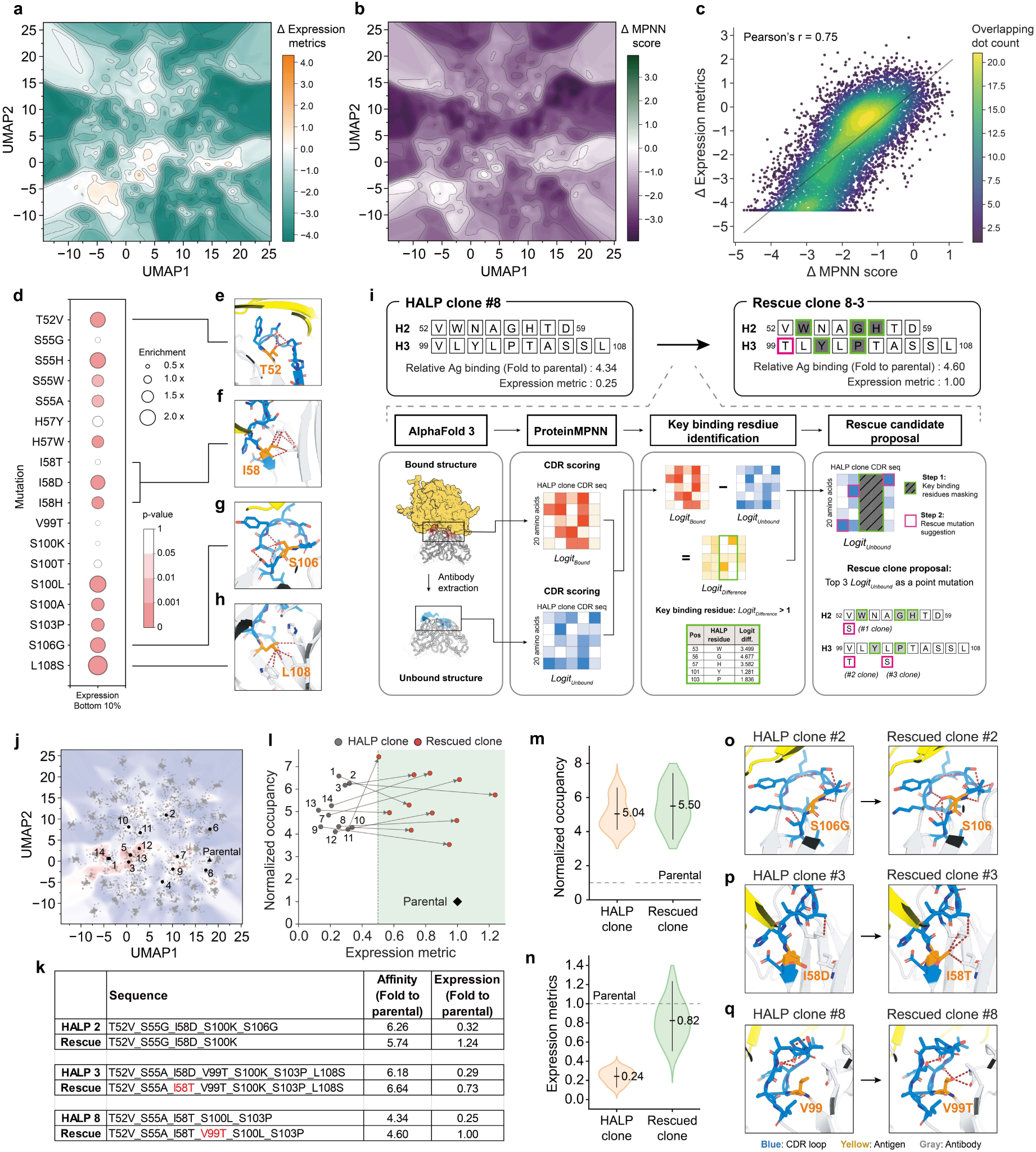
Computational profiling of expression liabilities and targeted rescue of high-affinity, low-production (HALP) variants. **a,b,** 2D contour landscapes illustrating the deviation (Delta) of (**a**) the experimental expression metric and (**b**) the ProteinMPNN compatibility score relative to the parental clone. Color gradients indicate variations from the parental baseline (white), with orange/dark green representing higher values and cyan/purple representing lower values. **c,** Correlation scatter plot between the expression metric and MPNN score (Pearson’s r = 0.75). The color scale reflects the overlapping dot count to visualize local data density. **d,** Bubble chart quantifying the enrichment of specific single mutations within the bottom 10% expression cohort. Bubble size denotes the enrichment ratio, while fill color indicates statistical significance (p-value). **e–h,** Structural snapshots of the parental complex highlighting the local interaction environments of key expression-limiting positions: (**e**) T52, (**f**) I58, (**g**) S106, and (**h**) L108. In all panels, the antigen is shown in yellow, the antibody framework in white, HCDR2 and HCDR3 in blue, and the target residue in orange. Key interactions are indicated by brick-colored dashed lines. **i,** Schematic representation of the computational rescue workflow for HALP clones. The pipeline illustrates the targeted recovery of expression while preserving antigen affinity, highlighted by the transition from HALP clone 8 to rescued clone 8-3. **j,** Projection of the 14 selected HALP clones onto the 2D UMAP affinity landscape. **k,** Summary table of three representative successful rescue strategies: reversion of a destabilizing mutation (HALP 2), substitution with an alternative mutation (HALP 3), and introduction of a novel compensatory mutation (HALP 8). **l,** Scatter plot tracking the two-dimensional fitness trajectory of the rescue process. Initial HALP clones (black) are connected via directional arrows to their successfully rescued counterparts (red), defined by achieving at least a two-fold productivity improvement relative to the initial HALP clone and reaching at least 50% of the parental level. **m,n,** Violin plots comparing the distributions and mean values of (**m**) relative occupancy and (**n**) the expression metric before and after the targeted computational rescue. **o–q,** Structural snapshots detailing the remodeled molecular interactions for the three successful rescue scenarios described in (**k**): (**o**) HALP clone 2, (**p**) HALP clone 3, and (**q**) HALP clone 8. The color scheme used in (**o**–**q**) follows the same convention as in (**e**–**h**).

We further investigated whether this predictive capability could be leveraged to re-engineer High-Affinity, Low-Productivity (HALP) clones (Fig. 4i). For each HALP sequence, we generated structural models of the antibody-antigen complex using AlphaFold3 and derived the antigen-free structure by removing the TNF-α coordinates, keeping the antibody backbone unchanged. This ensures that rescue mutations are evaluated within the binding-ready conformation, which is also expected to reduce the conformational entropy loss upon antigen binding. Both structures were then scored with ProteinMPNN to extract per-position logit values.

The rescue design proceeded in two steps. First, positions where the native amino acid’s logit score was substantially higher in the bound than the unbound conformation (logit difference > 1.0) were classified as binding key residues and excluded from mutation proposals. Second, the remaining positions were ranked by their native logit score in the unbound model, in ascending order, identifying sites of greatest sequence-structure mismatch in the antigen-free conformation. For each source sequence, we proposed a single substitution at each of the three lowest-scoring positions: the amino acid most strongly preferred by ProteinMPNN at that backbone position, yielding up to three candidate rescue mutations per HALP clone.

To test the generalizability of the expression rescue strategy across diverse antibody contexts, we selected HALP candidates spanning a broad range of CDR compositions from the variant library. Specifically, we applied k-means clustering on UMAP-projected sequence space and selected the highest-affinity representative from each cluster, yielding 12 source sequences with diverse CDR backgrounds. Two additional sequences were drawn from the high-affinity peak cluster identified in the fitness landscape, bringing the total to 14 (Fig. 4j). All 14 clones exhibited four- to seven-fold improvements in antigen affinity alongside an approximately five-fold reduction in cellular productivity relative to parental adalimumab (Fig. 4k,l and Supplementary Table 2).

To validate the strategy, we characterized the 42 single-residue rescue candidates across the 14 HALP sequences using the SPID workflow (Extended Data Fig. 8i-k). A successful rescue was defined as simultaneously retaining at least 80% of the source clone’s antigen affinity, achieving at least a two-fold productivity improvement relative to the source, and reaching at least 50% of parental adalimumab level. Remarkably, 11 of the 14 HALP sequences were successfully rescued, with at least one candidate meeting all three criteria (Fig. 4l-n and Supplementary Table 2). HALP clones 2, 3, and 8 particularly illustrate the mechanism (Fig. 4k): a single substitution improving interactions within the CDR or with the antibody framework yielded a 2.48- to 3.94-fold improvement in cellular productivity without compromising binding (Fig. 4o-q, Supplementary Note 2). These results demonstrate that structure-guided substitution of a single CDR residue can restore productivity in most HALP clones, largely preserving the binding properties that required extensive combinatorial optimization.

### *In vivo* therapeutic efficacy of landscape-navigated adalimumab variants

While the integration of our fitness landscapes with ProteinMPNN and AlphaFold3 enabled the structural rescue of compromised clones, these datasets also afford a more direct route for therapeutic optimization: navigating toward peak clusters where multiple fitness components are simultaneously enhanced. To assess whether this direct navigation delivers therapeutic advantages, we returned to clones 1207 and 1208, which were previously isolated from one such peak cluster (Fig. 3d). We first confirmed that when expressed and purified at preparative scales, clones 1207 and 1208 exhibited desirable developability profiles, including high cellular productivity and robust thermal stability, reproducing our initial assessment from the SPID workflow (Extended Data Fig. 9a-e). We then employed a TNF-α–responsive secreted embryonic alkaline phosphatase (SEAP) reporter assay (Fig. 5a). Both 1207 and 1208 clones exhibited stronger inhibition of TNF-α–induced signaling than parental adalimumab at matched concentrations, with IC₅₀ values of 15.3 pM and 26.2 pM, respectively, compared with 278.8 pM for adalimumab, demonstrating a substantial improvement in neutralizing efficacy (Extended Data Fig. 9f,g).

**Fig. 5.**
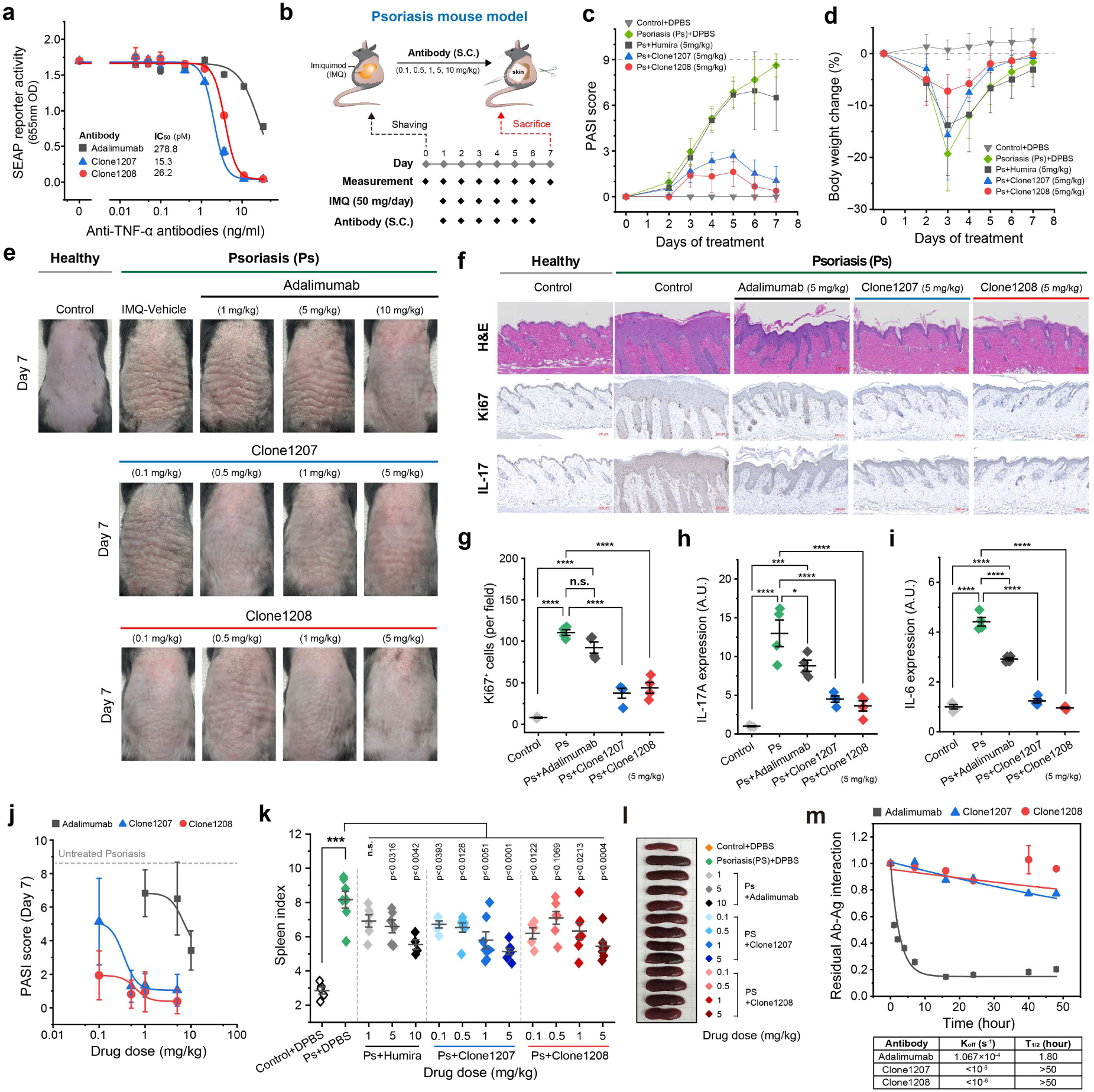
*In vivo* functional validation of SPID-engineered adalimumab variants for enhanced TNF-α neutralization. **a,** Functional inhibition of TNF-α signaling measured in an NF-κB responsive SEAP reporter assay using HEK293 cells. Concentration-response curves show improved neutralizing potency of engineered adalimumab variants clone1207 (IC_50_=15.3 pM) and clone1208 (IC_50_=26.2 pM) compared to Adalimumab (IC_50_=278.8 pM). **b,** Schematic of the imiquimod (IMQ)-induced psoriasis mouse model. **c,** Changes in daily Psoriasis Area and Severity Index (PASI) scores during treatment journey. **d,** Monitoring of body weight changes during treatment journey showing no overt toxicity. **e,** Representative macroscopic appearance of dorsal skin on day 7. **f,** Representative histological and immunohistochemical staining of dorsal skin lesions on day 7. The photographs show suppression of epidermal hyperplasia (H&E), reduced keratinocyte hyperproliferation (Ki67), and decreased inflammatory cytokine expression (IL-17A). The complete set of images for (**e**) and (**f**) is provided in **Extended Data** Fig. 10. **g,** Quantification of Ki67^+^ cells from tissue sections. **h,i,** Inflammatory cytokine levels within skin lesions: (**h**) IL-17A and (**i**) IL-6. **j,** Dose-response analysis of PASI scores at day 7 demonstrating dose-sparing efficacy of clones 1207 and 1208. **k,** Spleen index measured on day 7 as an indicator of systemic inflammation. **l**, Representative images of spleens collected on day 7. **m**, Comparison of residual Ab-Ag complex lifetimes between adalimumab variants and TNF-α using SPID.

We next utilized a psoriasis-like mouse model induced by imiquimod (IMQ), a Toll-like receptor 7 agonist. Mice received subcutaneous injections of either adalimumab or 1207/1208 according to the dosing schedule shown in Figure 5b. Clinical severity was evaluated daily using a composite psoriasis area severity index (PASI) that integrates keratinization, epidermal thickness and erythema. At an initial treatment dose of 5 mg/kg, 1207 and 1208 produced a rapid reduction in PASI scores beginning on day 2, with sustained improvement throughout the treatment period (Fig. 5c), whereas adalimumab showed comparatively lower efficacy at the same dose without overt body weight changes (Fig. 5d and Extended Data Fig. 9h). Macroscopic inspection of dorsal skin revealed prominent scaling and thickening in IMQ-treated mice (Fig. 5e and Extended Data Fig. 9i), corroborated by histological analysis of H&E-stained sections (Fig. 5f). Ki67 immunostaining demonstrated IMQ-induced keratinocyte hyperproliferation, and interleukin-17A (IL-17A) and interleukin-6 (IL-6) expressions within lesional skin confirmed heightened inflammatory signaling (Fig. 5g-i). Across these readouts, 1207 and 1208 consistently outperformed adalimumab, indicating an enhanced capacity to attenuate TNF-α–driven inflammatory responses.

We systematically varied treatment dose to define the therapeutic windows of the three antibodies. Adalimumab required a dose of 10 mg/kg to elicit a robust therapeutic response. By contrast, 1207 and 1208 achieved comparable efficacy at substantially lower doses, as low as 0.5 mg/kg and 0.1 mg/kg, respectively, which corresponded to 20- to 100-fold dose reductions relative to parental adalimumab (Fig. 5j and Extended Data Fig. 9j). At these reduced dosing levels, mice treated with 1207 or 1208 also exhibited significantly attenuated splenomegaly, indicative of reduced IMQ-induced systemic inflammation (Fig. 5k,l). These findings suggest that 1207 and 1208 confer immunomodulatory effects extending beyond local skin improvement, mitigating systemic inflammatory burden in this psoriasis-like model.

The magnitude of the *in vivo* potency gain (20- to 100-fold) was striking given the more modest enhancement in equilibrium antigen occupancy (several-fold) (Fig. 5j,k versus Extended Data Fig. 9e), prompting us to investigate whether an additional biophysical parameter underlies the therapeutic advantage. Consistent with this hypothesis, direct tracking of the lifetime of adalimumab–TNF-α complexes using the SPID imaging system revealed that 1207 and 1208 displayed minimal complex dissociation over an observation period of 50 hours, whereas the parental adalimumab exhibited a complex half-life of approximately 1.8 hours (Fig. 5m). This remarkable >25-fold extension in complex lifetime aligns with the magnitude of the observed *in vivo* potency gain, indicating that target residence time, rather than mere equilibrium affinity, mechanistically underlies the therapeutic advantage of clones 1207 and 1208. Together, these findings establish that landscape-scale navigation can yield antibody variants with superior *in vivo* efficacy through the optimization of diverse biophysical properties, such as complex lifetime, extending beyond conventional affinity optimization.

## Discussion

Antibodies have long been viewed as binders whose CDR geometry is shaped primarily by antigen engagement, a framework that has guided decades of antibody engineering centered around binding affinity^23,100^. Our SPID-generated landscape-scale data brings a complementary constraint into focus. Within the native adalimumab backbone, the productivity consequences of thousands of CDR sequence variants were accurately predicted by ProteinMPNN, a generic inverse-folding model not specifically trained for antibodies^48^. This indicates that CDR residues, despite the conformational diversity intrinsic to CDRs and particularly pronounced in HCDR3^101,102^, contribute to cellular productivity through the same sequence-structure compatibility principles that apply to folded proteins more broadly. Where reliable structural models exist, our observation has a direct implication: productivity liabilities introduced during affinity maturation, humanization, or CDR grafting can be interpreted, scored, and addressed through the same inverse-folding framework that guides general protein engineering^103,104^. Moreover, when applying this scoring scheme to cases where the structural template for the antibody was generated using structure prediction engines such as AlphaFold3^49^, we confirmed the validity of the ProteinMPNN predictions, albeit with slightly increased error rates (data not shown). Together, our findings suggest that CDR loops adopt defined structural folds, governed by the same biophysical principles as globally folded proteins, long before any antigen encounter, and that mutations destabilizing these nascent folds impair cellular expression.

The orthogonality of affinity and productivity determinants, first suggested by our single-residue DMS across ten therapeutic antibodies and made concrete in the combinatorial landscape of adalimumab, has practical as well as mechanistic consequences. Because the fitness cliffs of the two properties do not coincide, windows open within the sequence space where one trait can be improved without sacrificing the other. The rescue of HALP clones illustrates this principle in concrete form: a single substitution that restores nascent structural compatibility is often sufficient to recover cellular productivity without eroding the binding properties assembled through extensive combinatorial screening. What makes this outcome notable is the asymmetry it exposes. While hard-won affinity-enhancing mutations are fragile and difficult to maintain during downstream optimization, productivity defects are frequently locally correctable when guided by precise structural readouts^30,91^. This asymmetry has a direct consequence for antibody engineering pipelines: affinity and productivity screening no longer need to be strictly coupled at the selection stage^21^. High-affinity candidates burdened by productivity liabilities, historically discarded as dead ends, can now be systemically salvaged. Finally, this asymmetry clearly delineates the current frontier of AI-driven antibody design. Although state-of-the-art AI models can reliably predict the nascent CDR structural folds and assess their sequence-structure compatibility, fully unlocking de novo antibody design will still require mastering the complex conformational plasticity that drives active, high-affinity antigen engagement.

Of note, these findings and conclusions essentially rest on a technical shift enabled by the SPID workflow. Landscape-scale profiling at single-molecule precision maps the complex topography of multiple fitness landscapes with unprecedented resolution—exposing an extreme ruggedness and non-navigability in both affinity and productivity spaces that conventional screens have been largely blind to^105^. Because our combinatorial library was built exclusively from functional variants pre-filtered via DMS, this rugged topology serves as direct evidence of pervasive negative epistasis among otherwise beneficial mutations. Nevertheless, rare instances of positive epistasis did occur, generating the observed peak clusters and the global top 1% of clones. As most of these elite clones harbored four to seven simultaneous mutations, a critical challenge emerges: how to efficiently navigate these rugged landscapes without resorting to exhaustive combinatorial screening, which scales exponentially. While the stepwise accumulation of beneficial mutations is a standard evolutionary trajectory, our pathway analyses for clones 1207 and 1208 demonstrated the futility of this approach, revealing a complete absence of pathways offering monotonic fitness gain.

Instead, by taking advantage of the remarkable flexibility in library production and characterization afforded by the SPID workflow, we demonstrate pattern recognition for epistatic interactions at smaller scales enables highly efficient sampling of the rugged fitness landscapes. For instance, employing machine learning to capture the simplest epistatic interactions—pairwise mutations^91,99^—drastically condensed the search space by eliminating variants that failed at the double-mutant stage^25,27^. Furthermore, a modular assembly strategy that recombines top-performing intra-CDR variants effectively prioritizes local epistatic interactions within CDRs over rarer inter-CDR interactions. This modular approach proved remarkably efficient, successfully recovering 70% of the global top 1% clones by screening only the top 20% of individual CDR combinations. Importantly, these machine-learning and modular strategies are not mutually exclusive; synergizing these approaches to map epistatic networks enables a streamlined navigation of immense combinatorial spaces. Together, these findings position SPID-driven landscape-scale profiling not merely as an incremental advance in screening throughput and precision, but as a transformative toolset for interrogating antibody sequence space, —yielding the dense, quantitative ground-truth datasets poised to propel the evolution of AI-guided antibody design.

## Materials and Methods

### Antibodies and drugs

Biotin anti-RFP antibody (PRO50-00064-00; Proteina), anti-human IgG Fcγ with biotin conjugation (109-066-098; Jackson Immunoresearch), and anti-human IgG (Fab’)2 with biotin conjugation (ab98533; Abcam) were used to immobilize the scFabs for SPID as well as used to detect the scFabs by western blotting. Trastuzumab (HY-P9907; MedChemExpress) and Adalimumab (HY-P9908; MedChemExpress) were used to competitive binding assay for SPID. Human anti-adalimumab antibodies (HCD203, HCD204; Bio-Rad Antibodies) were used to an AIA interaction assessment for SPID. HRP-linked streptavidin (N100; Thermo fisher Scientific) was used to label and detect the protein levels for western blotting. Anti-Ki67 (MA5-14520; Invitrogen), anti-IL6 (P620; Invitrogen), anti-IL17A (ab79056; Abcam), anti-GSDMD (10137; Cell Signaling Technology), anti-NLRP3 (AG-20B-0014-C100; AdipoGen), and anti-ASC (AG-25B-0006-C100; AdipoGen) antibodies were used to immunohistochemistry.

The antibody drugs (adalimumab, anti-TNF-α and anti-GIPR antibodies) antibodies used for the SPID assay development, SEAP assay, and animal experiments were custom-produced and purified by Y-Biologics (South Korea).

### Animals

Animal experiments were approved by the Institutional Animal Care and Use Committee of Seoul National University (SNU; IACUC No. SNU-250416-1). Male C57BL/6 wild-type mice (6 weeks old) were purchased from JA Bio Inc. (South Korea). Mice were housed under controlled conditions (23–25°C, 45–65% humidity) with a 12-hour light/dark cycle. All procedures complied with relevant ethical regulations.

### Cell culture and collection

HEK293T cells were purchased from American Type Culture Collection (ATCC). HEK-Blue™ TNF-α cells were purchased from InvivoGen. HEK293T cells were cultured in DMEM (D6429; Sigma Aldrich) supplemented with 10% (v/v) FBS (26140-079; Gibco) and 100 μg/ml penicillin/streptomycin (15140-122; Gibco). SK-BR-3 cells were cultured in RPMI1640 (R8758; Sigma Aldrich) supplemented with 10% (v/v) FBS and 100 μg/ml penicillin/streptomycin. The cells were incubated in a humidified incubator at 37 °C, 5% CO_2_. HEK-Blue™ TNF-α cells were cultured in DMEM supplemented with 10% (v/v) FBS, 100 μg/ml penicillin/streptomycin, and 100 µg/ml normocin (ant-nr-2; InvivoGen).

For cell collection, cultured cells were rinsed with cold DPBS (D8537; Sigma Aldrich) and then were collected with the scraper in 1 ml of cold DPBS. The collected cells were centrifuged at 500 g for 5 minutes at 4 °C, and the supernatants were discarded. The cell pellets were stored at - 80 °C by snap-freezing with liquid nitrogen.

### Recombinant antibody expression

The variable regions of scFab antibodies were cloned into *pCMV* mammalian expression vectors with the constant domains of a common constant regions obtained from human IgG_1_ (CH: ASTKGPSVFPLAPSSKSTSGGTAALGCLVKDYFPEPVTV-SWNSGALTSGVHTFPAVLQSSGLYSLSSVVTVPSSSLGTQTYICNVNHKPSNTKVDKKV EP, LH: RTVAAPSVFIFPPSDEQLKSGTASVVCLLNNFYPREAKVQWKVDNALQSGNSQ-ESVTEQDSKDSTYSLSSTLTLSKADYEKHKVYACEVTHQG-LSSPVTKSFNRGEC). For recombinant IgG antibodies, the variable regions were similarly cloned into human IgG_1_ constant domains. All cDNAs of variable regions of recombinant antibodies (Trastuzumab, Pertuzumab, Atezolizumab, Obinutuzumab, Elotuzumab, Adalimumab, Ipilimumab, Durvalumab, Avelumab, Ustekinumab, Infliximab, Rituximab, Pembrolizumab, Nivolumab, Panitumumab, Cetuximab, Camrelizumab, Cemiplimab, Serplulimab, Dostarlimab, Tislelizumab, Bevacizumab, Ranibizumab, Eculizumab, Etesevimab, Dupilumab, Tixagevimab, DISC0100 (anti-IL15), UFKA-20 (anti-IL2), M116 (anti-CCL17), CNTO888 (anti-CCL2), E10 (anti-CXCL13), LY3041658 (anti-CXCL3), MOR04302 (anti-GM-CSF), MOR8457 (anti-PDGF-BB), COV89-22 (anti-COVID19 SH2), and anti-GIPR, and H7 (anti-TfR1)) were synthesized by Macrogen (South Korea). The 18 different restriction enzyme recognition sites were removed in the *pCMV* mammalian expression vectors, therefore all cDNAs were cloned by Gibson assembly (E2611; New England Biolabs).

In the scFab format, the heavy chain and light chain domains were linked with 60-length of polypeptide linkers (GGSSGSGSGSTGTSSSGTGTSAGTTGTSASTSGSGSGGGGGSGGGGSAGGTATAGASSGS). For the recombinant IgG format, the heavy and light chains were encoded within a single vector, interconnected by a Furin recognition site (RRKR) and a T2A self-cleaving peptide sequence (EGRGSLLTCGDVEENPGP). The signal peptide of the human immunoglobulin heavy chain (MDWTWRILFLVAAATGAHS) and the mScarlet protein were fused to the N-terminus and the C-terminus of scFab protein, respectively. Regarding the IgG constructs, the signal peptides of the human immunoglobulin heavy chain (MDWTWRILFLVAAATGAHS) and light chain (MLPSQLIGFLLLWVPASRG) were fused to the N-terminus of each respective chain, and the mScarlet protein was fused to the C-terminus of the heavy chain.

The resulting plasmids were introduced into HEK293T cells through transient transfection to overexpress the mScarlet-labeled antibody proteins. 5×10^5^ of HEK293T cells were seeded into 6-well cell culture plates in a day before transfection. 2 μg of plasmid DNA was mixed with 1:3 (w/w) ratio of PolyJet (SL100688; SignaGen) in 0.1 ml DMEM, and the mixture was introduced into the seeded HEK293T cells. The cell culture media containing antibodies were collected 2 days after transfection. The cell debris were eliminated by centrifugation at 500 g for 5 minutes at 4 °C, and the supernatants were stored at – 80 °C after snap-freezing.

### Antigen protein expression

The cDNAs of antigen proteins (HER2, EGFR, PD1, PDL1, CTLA4, CD20, SLAMF7, VEGFA_165_, TNF-α(77-233), C5(819-932), IL12b, IL2, IL15, IL4Rα, CCL17, CCL2, CXCL13, CXCL3, GM-CSF, PDGF-BB, GIPR, and TFR1) were isolated from their respective human cDNA. The cDNAs RBD and SH2 domain of COVID19 were synthesized by the Macrogen (South Korea). All cDNAs were cloned into *pCMV* vectors with eGFP sequences on the C-terminus to generate eGFP-labeled antigen constructs, except for TFR1, which was fused with eGFP at the N-terminus. The resulting plasmids were introduced into HEK293T cells through transient transfection to overexpress the eGFP-labeled antigen proteins. 3×10^6^ of HEK293T cells were seeded into 90mm^2^ cell culture dishes in a day before transfection. 10 μg of plasmid DNA was mixed with 1:3 (w/w) ratio of PolyJet in 1 ml DMEM, and the mixture was introduced into the seeded HEK293T cells. For the intracellular expressing proteins, the cells were collected 24 hours after transfection and stored at -80 °C. While the secretory proteins, the cell culture media containing the proteins were collected 24 hours after transfection. The cell debris were eliminated by centrifugation at 500 g for 5 minutes at 4 °C, and the supernatants were stored at – 80 °C after snap-freezing.

### Cell lysis for antigen preparation

The collected cell pellets expressing antigen proteins (HER2-eGFP, PDL1-eGFP, CTLA4-eGFP, CD20-eGFP, TNF-α-eGFP, PD1-eGFP, and EGFR-eGFP) were resuspended for cell lysis with the TX100 lysis buffer (1% (v/v) Triton-X100, 1% (v/v) protease inhibitor (PRO50-00017-00; Proteina), 1% (v/v) phosphatase inhibitor cocktail 2 (PRO50-00018-00; Proteina), 1% (v/v) phosphatase inhibitor cocktail 3 (PRO50-00019-00; Proteina) in lysis base buffer (PRO50-00005-00; Proteina)). For the cell pellets expressing antigen proteins (CD20-eGFP, SLAMF7-eGFP, IL4Rα-eGFP, GIPR-eGFP), the glyco-diosgenin (GDN) lysis buffer (1% (w/v) GDN (GDN101; Anatrace) in lysis base buffer) was used to cell lysis. The eGFP-TFR1 expressing cells were lysed with the lauryl maltose neopentyl glycol (LMNG) lysis buffer (1% (w/v) LMNG (NG310; Anatrace) in lysis base buffer).

After resuspending the cell pellets with the designated lysis buffer, the cell suspensions were incubated for 30 minutes at 4 °C. The supernatants were isolated after 15,000 g centrifugation for 10 minutes at 4 °C, and the concentration of the fluorescence proteins containing in supernatant was measured. The supernatants were stored at -80 °C followed by snap-freezing.

### Western blotting

The loading concentration of each scFab sample was quantified based on the mScarlet concentration. Before the SDS-PAGE, all the scFab samples were heated at 95 °C for 10 minutes with SDS containing loading dye (EBA-1052; ELPIS Biotech). The denatured samples were resolved with 12% SDS-PAGE gels, and then transferred to PVDF membranes included in iBlot Transfer System (IB401001; Thermo Fisher Scientific). The membranes were blocked in 5% (v/w) skim milk in TBST buffer (20 mM Tris pH 7.6, 150 mM NaCl, 0.1% (w/v) Tween 20) for 1 hour at room temperature, and immunoblotted with aforementioned antibodies diluted 1:2,000 in TBST buffer with 1% (w/v) BSA overnight at 4 °C. The membranes were then blotted with aforementioned HRP-linked streptavidins diluted 1:4,000 in TBST buffer with 1% (w/v) BSA. The protein bands were detected with an ImageQuant LAS 4000 mini luminescent analyzer (ImageQuant LAS 4000 mini; Cytiba) by using an ECL (W3653-020; GenDEPOT) chemi-illumination system.

### High-throughput in vitro CDR editing

To construct modular vectors for CDR substitutions, the modular vectors were engineered by replacing the native CDR or variable region (either V_H_ or V_L_) with a synthetic insert flanked by outward-facing restriction enzyme recognition sites. To prevent non-functional or leaky expression, five stop codons were inserted between the restriction sites, ensuring that only correctly assembled constructs would result in recombinant antibody expression.

For single-CDR substitution, the modular plasmid vectors were digested with pre-designated restriction enzymes at 37 °C for 2 hours to discard the CDR encoded regions. After digestion, the vectors were resolved on a 0.8% agarose gel, and the digested vectors were selectively purified with agarose gel purification system (QIAGEN). Single-stranded DNAs (ssDNAs) encoding the CDR sequences were synthesized by Macrogen (South Korea) or Integrated DNA Technologies (IDT, USA). The initially synthesized forward and reverse ssDNAs were rehydrated respectively, and mixed with 1:1 ratio to achieve a final concentration of 50 μM in annealing buffer (10 mM Tris pH 7.6, 50 mM NaCl, 1 mM EDTA). The ssDNA mixture was annealed via heating followed by progressive cooling (95 °C, 5 minutes; 55 °C, 30 minutes; 25°C, 5 minutes; 15°C, 5 minutes; ramp rate was 0.1 °C/s) to obtain 50 μM of complementary double-stranded DNAs (dsDNAs). The integrity of the annealed dsDNA fragments was confirmed by 18% PAGE gel and detected after SYBR Gold staining (S11494; Thermo Fisher Scientific). The digested vectors and the annealed dsDNAs were mixed at a molar ratio of 1:30, and ligated using T4 DNA ligases (M0202; New England Biolabs) in T4 DNA Ligase Reaction Buffer (B0202; New England Biolabs) for 2 hours at 16 °C. For multiple CDR substitutions, 400ng of modular vector was combined with equimolar amounts of dsDNA fragments encoding CDR variants and framework regions using Golden Gate enzyme mix (E1601; New England Biolabs) in T4 ligase buffer. The DNA fragments were designed with compatible sticky ends and prepared by annealing complementary ssDNAs. When targeting multiple CDRs simultaneously, T4 polynucleotide kinase (M0201; New England Biolabs) was added for fragment phosphorylation. The molar ratio of vector to total inserts was maintained at 1:30, and reactions were carried out with the following thermal protocol (37 °C, 2 hrs; 65 °C, 30min; 37 °C, 1hr).

The 100 ng of resulting ligated plasmids were introduced into 1.8×10^4^ HEK293T cells seeded in a 96-well cell culture plate through transient transfection. After 2 days of expression, the cell culture media containing recombinant antibodies were collected.

### SPID

SPID was performed using the Pi-Chip (PC04000, PC38400; Proteina) and Pi-View (PRS51-00003-00; Proteina) system. Bind Solution (PRO05-00006-00; Proteina) was diluted with 1:100 by T buffer and loaded into each reaction chamber of the Pi-Chip. After washing the imaging chip with T buffer, biotin anti-RFP antibodies were loaded to each reaction chamber with 1:1,000. Then, cell culture media containing recombinant antibodies were loaded into each reaction chamber to immobilize the antibodies on the chip surface with 1:500 (typically, 20 pM) for 30 minutes. After washing the reaction chambers with T buffer, the cell extracts containing eGFP-labeled antigens were loaded into each reaction chamber with designated concentration. After being incubated for 2 hours to reach the binding equilibrium, the chip was moved on to Pi-View system to acquire fluorescence images.

We employed the methods outlined in previous study for single-molecule fluorescence imaging by using SPID^43^. In detail, single-molecule fluorescence imaging was performed with two different excitation lasers for 488 nm and 561 nm wavelengths. The emitted photons were collected by sCMOS camera in Pi-View system and converted to the single-molecule fluorescence images. From the obtained fluorescence images, the single-molecule spots were determined by Pi-Analyzer software (Proteina, South Korea). The number of surface-immobilized antibodies (RFP signals) and the number of antibody-antigen complex (GFP signals) were converted to fractional occupancy, and then the dissociation constants (*K*_D_s) were calculated by the following equation.

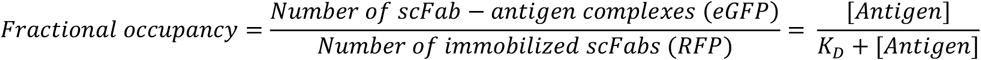

### SPID-based thermal stability assay

Antibody stability was evaluated by applying heat treatment using a thermal cycler directly to unpurified cell culture supernatants containing antibodies. The supernatants were heated at a designated temperature (e.g., 60 °C) for 10 minutes and immediately cooled to 4 °C. Subsequently, the fractional occupancies for antigen were measured using SPID. The relative thermal stability was calculated as the ratio of the fractional occupancies from heat-treated antibodies to those from untreated antibodies. Thermal stability was determined as the residual binding capacity, defined as the ratio of the fractional occupancy after heat treatment to that before treatment (Extended Data Fig. 3e). Specifically, the relative Ab-Ag binding represents the retention of binding occupancy under thermal stress. For heatmap analysis, the normalized Ab-Ag binding was calculated by dividing the relative Ab-Ag binding value of each variant (at 60 °C) by the corresponding value of the parental antibody.

### DSF

To determine the melting temperature (T_m_) of purified antibodies using a conventional method, DSF was performed. Purified antibody samples were diluted to a final concentration of 0.2 mg/mL in a phosphate-buffered saline (PBS, pH 7.4) solution. SYPRO Orange Protein Gel Stain (S6650; Invitrogen) was added to the protein solution at a 5X final concentration from a 5,000X stock. The assay was performed in a 96-well PCR plate, with each well containing a 20 uL of final volume. The plate was heated in a real-time PCR system (QuantStudio 3; Applied Biosystems) from 25 °C to 95 °C at a ramp rate 1 °C /min. Fluorescence intensity was monitored continuously using the TAMRA channel. The raw fluorescence data was plotted against temperature, and the T_m_ was determined by identifying the peak of the first derivative of the melting curve using the DSFworld^106^.

### Physicochemical vectorization and densMAP projection

To visualize the macroscopic topography of the combinatorial sequence library (∼9,600 variants), the sequence space was parameterized based on physicochemical properties rather than one-hot encoding. For the nine targeted mutational residues within the adalimumab HCDR2 and HCDR3 loops (positions 52, 55, 57, 58, 99, 100, 103, 106, and 108), each amino acid substitution was encoded as a three-dimensional vector representing hydrophobicity, side-chain volume, and isoelectric point (PI) (**Extended Data Fig. 7**). To precisely capture the structural and biochemical deviation from the parental clone, parental residues were zero-centered as the origin ([0, 0, 0]) in the property space. Consequently, each variant was represented as a continuous 27-dimensional feature vector. Following Z-score standardization, these high-dimensional vectors were projected onto a two-dimensional embedding using the Uniform Manifold Approximation and Projection (UMAP) algorithm. To preserve both the local topological structure and the global density distribution of the mutational space—thereby preventing visual distortion of dense variant populations—the density-preserving variant of UMAP (densMAP) was strictly employed. Hyperparameters were empirically optimized with the number of nearest neighbors set to 50, the effective minimum distance to 0.1, and the effective scale of embedded points to 2.0.

### Topographical fitness landscape construction

Continuous three-dimensional fitness landscapes were constructed using the 2D densMAP coordinates as the basal plane (x, y). Experimental functional metrics (log-normalized relative fractional occupancy and expression metrics) were mapped onto the z-axis to establish robust sequence-to-function relationships. This approach enabled the visual and quantitative translation of the mutational sequence space into a macroscopic topographical landscape. Based on the spatial continuity and variation of these fitness metrics, the geometric architecture of the combinatorial library was characterized into distinct topographical features, allowing for a comprehensive assessment of functional adaptation driven by physicochemical properties.

### Combinatorial search space reduction and optimal sequence identification

To circumvent the prohibitive cost and time associated with exhaustive screening of massive combinatorial libraries, two distinct strategies were employed to compress the search space while recovering the global top 1% optimal variants.

For the modular assembly of HCDR2 and HCDR3, we firstly leveraged the structural rationale that mutations within the loops contribute independently to the overall functional fitness. By pre-evaluating the individual loops and selecting only top-tier modules, the search space was effectively truncated. These high-ranking modules were subsequently subjected to combinatorial pairing, drastically reducing the total number of variants required for synthesis while maintaining a high probability of identifying global optima.

A second strategy utilized a machine learning (ML)-guided positive design approach to infer optimal mutational combinations from sparse initial datasets. The sequence space was compressed by training a model exclusively on empirical fitness data (log-normalized fractional occupancy and expression metrics) derived from low-order mutants, specifically variants harboring up to double or triple amino acid substitutions. Sequences were parameterized for the ML model via one-hot encoding (MultiLabelBinarizer), transforming the presence or absence of individual mutations into a multidimensional binary matrix. A Ridge linear regression model (regularization parameter α= 1.0) was then fitted to these binary feature vectors and their corresponding experimental fitness scores. This linear modeling allowed for the deconvolution of the sequence-function relationship, where the resulting regression coefficients represented the independent, additive weights of each single amino acid mutation. Mutations exhibiting negative coefficients (deleterious effects) were systematically eliminated from the design space. A computationally focused library was then constructed by imposing a strict combinatorial constraint: only sequences composed entirely of mutations with positive additive weights (positive design) were permitted. This data-driven filtering approach enabled the recovery of global top 1% variants with a significantly minimized synthesis budget.

### ProteinMPNN probability-based scoring for landscape analysis

Scoring was performed using ProteinMPNN (proteinmpnn_v_48_020.pt) in single-residue scoring mode, specified by the flags --single_aa_score 1 and --use_sequence 1. A batch size of 10 (--batch_size 10) was used to aggregate scores across multiple independent forward passes, improving the stability of the estimates. In single-residue scoring mode, the model computes the conditional probability of the native amino acid at each position given the backbone structure and the surrounding sequence context, *p(AAi | backbone, AA_≠i_)*. A high probability at a position indicates that the residue is well accommodated by the local backbone geometry; a low probability indicates incompatibility between the amino acid identity and the structure it must adopt.

Per-position probabilities were aggregated into a *compatibility score* representing the mean negative log-probability of the native sequence across all evaluated residues:

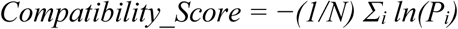

where *P_i_* is the mean softmax probability of the native amino acid at position *i* and *N* is the number of evaluated residues. A lower compatibility score indicates greater sequence-structure compatibility. To compare this metric directly with experimental expression data, where higher values reflect better expression, we defined a normalized compatibility score *Y* for each variant relative to adalimumab:

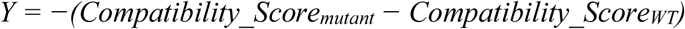

Positive Y values indicate improved structural compatibility relative to adalimumab.

### AlphaFold3 modeling

Structural models of the antibody-antigen complex were generated using AlphaFold3 for each of the 14 HALP source sequences, sampling 5 random seeds per sequence to produce 25 structural models per candidate. The model with the highest pLDDT across HCDR2 and HCDR3 was selected as the representative bound-state structure. The antigen-free unbound structure used for rescue scoring was then derived by removing the TNF-α coordinates from the selected bound-state model, keeping the antibody backbone unchanged. Both the bound and unbound structures were subsequently used for ProteinMPNN scoring.

### ProteinMPNN logit-based scoring for expression rescue design

For the rescue analysis, we modified the default ProteinMPNN scoring script (score.py) to additionally extract and save the mean (mean_logits) and standard deviation (std_logits) of the raw pre-softmax logits at each position across all 20 amino acids. This modification does not alter the model weights or inference procedure. Scoring was performed on both the bound and unbound antibody structures (derived as described above), using the aforementioned model weights and scoring mode.

Logits rather than probabilities were used for two operations in the rescue analysis. First, the bound-minus-unbound logit difference is used to classify binding key residues, because it provides a position-resolved measure of how much the antigen context selectively favors the native amino acid identity, with greater dynamic range than the softmax-compressed probability difference. Second, positions are ranked for expression rescue by the native amino acid’s logit score in the unbound model, which more sensitively resolves the degree of sequence-structure mismatch than ranking by probability alone.

### Rescue mutation design: Binding key residues classification and substitution proposals

#### 1) Binding key residue classification

For each position *i* in HCDR2 and HCDR3, the bound-minus-unbound logit difference for the native amino acid was computed as:

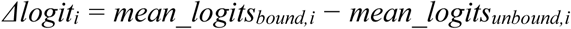

Positions satisfying *Δlogit_i_ > 1* were classified as binding key residues and excluded from mutation proposals. This threshold reflects positions where the native amino acid is significantly more favored in the presence of the antigen than in its absence, indicating that the residue identity is constrained by antigen contact.

#### 2) Ranking of expression-limiting positions and mutation proposals

Among residues that are not classified as binding key residues, each was ranked by the native amino acid’s mean_logits value from the unbound model, in ascending order. For each source sequence, the three positions with the lowest native logit scores were selected. At each position, the proposed rescue substitution was the amino acid with the highest mean_logits value in the unbound model at that position. This yielded up to three candidate rescue mutations per source sequence.

### Selection of candidates for expression rescue

#### 1) Filtering criteria

The initial dataset of 9,517 variants was filtered using the following criteria:

- Primary filter: Normalized_Occupancy > 4 and Normalized_Expression < 0.5.
- Error-corrected filter: (Normalized_Occupancy − error) > 3 and (Normalized_Expression + error) < 0.5.

The error term for each measurement was derived from replicate experiments and represents the standard deviation of the normalized value. This additional filter ensures that selected sequences robustly satisfy the high-affinity, low-expression criterion rather than qualifying only at the boundary of measurement uncertainty.

#### 2) Diversity-aware selection

The filtered subset was projected into a two-dimensional UMAP space and clustered using k-means with k = 12. The variant with the highest Normalized_Occupancy within each cluster was selected as the representative source sequence, yielding 12 sequences that are well distributed across the UMAP projection space and represent diverse HCDR2 and HCDR3 backgrounds. Two additional sequences were selected from the high-affinity peak cluster identified in the fitness landscape, bringing the total to 14 source sequences.

### IMQ-induced psoriasis model

Male C57BL/6 mice (6 weeks old) were acclimated for ≥6 days in a specific pathogen-free facility with ad libitum access to food and water. To induce psoriasis, mice were shaved and depilated on a 2 × 3 cm area of dorsal skin on day 0. Adalimumab (1, 5, or 10 mg kg⁻¹), clone1207, and clone1208 (0.1, 0.5, 1, or 5 mg kg⁻¹) were administered subcutaneously 30 min prior to imiquimod (IMQ) application. IMQ cream (50 mg day⁻¹) (5% Aldara cream; IMQ Dong-A ST) was applied topically to the exposed skin for 7 consecutive days. Body weight and clinical severity scores were recorded daily. On day 7, dorsal skin was photographed, and tissue samples were collected.

### PASI scoring and body weight change

Psoriasis severity was assessed daily using a modified PASI scoring system adapted for murine models. Skin thickness is measured with electric calipers (Digital Thickness calipers 0-150mm) (M12.500181; Mitutoty) across a fold of skin. Erythema, scaling, and thickness of the dorsal skin were each scored on a scale from 0 to 3 (0 = none, 1 = mild, 2 = moderate, 3 = severe), yielding a cumulative score ranging from 0 to 9 per mouse. Scoring was performed by blind observers under consistent lighting conditions. Body weight was measured daily at consistent time points throughout the 7-day imiquimod treatment period using a calibrated digital scale. Body weight (BW) data were normalized to baseline values recorded on day 0 and expressed as percentage change. Body weight (BW) change was calculated as follows: BW change (%) = [(Weightday_n_ – Weightday_1_) / Weightday_1_] × 100%. Graphs were generated using GraphPad Prism (v10.0), and data are presented as mean ± standard deviation (SD). Statistical significance was determined using repeated-measures analysis of variance (ANOVA), followed by Šídák’s multiple comparisons test.

### Measurement of back skin and epidermal Thickness

Back skin thickness was assessed using an Electric vernier caliper on experimental every day. During measurement, one operator restrained the mouse to expose the dorsal skin, while a second operator randomly selected three sites for evaluation. The thickness of the double-layered skin was recorded at each site. For each mouse, the change in skin thickness was calculated as the mean thickness on the designated day minus the mean thickness on day 1.

### Spleen index

The isolated spleen was washed with PBS to remove blood and impurities. After carefully removing the surrounding tissues, the spleen was weighed. Spleen index was determined using the formula: spleen weight (mg) / body weight (g), as an indicator of splenic hypertrophy.

### Histology and immunohistochemistry

Skin tissues were harvested from experimental sites and fixed in 10% neutral-buffered formalin. Fixed samples were dehydrated through a graded ethanol series, cleared in xylene (Sigma), and embedded in paraffin. Paraffin blocks were sectioned at 5 μm thickness and mounted onto glass slides.

For immunohistochemical analysis, paraffin-embedded sections were deparaffinized in xylene and rehydrated through graded ethanol (100%, 95%, 80%, and 70%). Antigen retrieval was conducted by microwave heating in citrate buffer (10 mM sodium citrate, 0.05% Tween 20, pH 6.0). Endogenous peroxidase activity was blocked with 3% hydrogen peroxide for 10 min. Non-specific binding was blocked by incubating sections with 10% normal goat serum (Invitrogen, Thermo Fisher Scientific) for 30 minutes at room temperature. Primary antibodies (TNF-α, IL-6, IL-17, and Ki-67) were applied with 1:100 dilution and incubated overnight at 4 °C. Following PBS washes, sections were incubated with HRP-conjugated secondary antibodies (Proteintech) for 2 hours at room temperature and washed three times with PBS. Signal detection was performed using DAB substrate solution (Vector Laboratories). Counterstaining was performed with hematoxylin. Dehydration was carried out using ascending ethanol concentrations (70%, 80%, 95%, and 100%), followed by clearing in xylene for 5 minutes, and was mounted using Fisher Chemical™ Permount™ Mounting Medium (Fisher scientific). Microscopic analysis was conducted using the Axio Scan Z1 Digital Slide Scanner (Carl Zeiss, Germany).

### SEAP reporter assay

HEK-Blue™ TNF-α cells are stably expressing a SEAP reporter under the control of an NF-κB/AP-1-responsive promoter. Activation of TNF-α signaling of induces NF-κB/AP-1 activity and subsequent SEAP secretion, which is quantified colorimetrically using Quanti-Blue™ solution (InvivoGen) according to the manufacturer’s instructions. All experiments were performed with cells maintained below passage 20.

Antibody variants (clone1207, 1208) and adalimumab (reference control) were tested at final in-well concentrations ranging from 33 ng/ml to 0.6 ng/ml in the presence of 1 ng/ml recombinant human TNF-α (300-01A; PeproTech). Recombinant human IL-1β (1 ng/ml) was included as a negative control. Antibodies and cytokines were prepared with 0.1% bovine serum albumin (BSA) (0.2-µm filtered) and dispensed into 96-well plates before adding HEK-Blue™ TNF-α cells (2 × 10⁴ cells in medium without selection antibiotics). After 24-hour incubation at 37 °C, 5% CO₂, 20 µl of conditioned medium was transferred to a fresh plate containing 180 µl Quanti-Blue™ solution (rep-qbs; InvivoGen). Plates were incubated for 1 hour at 37 °C, and SEAP activity was determined colorimetrically at 655 nm using a microplate reader.

## Code availability

The custom MATLAB and Python codes used for all analysis are available on GitHub at https://github.com/tyyoonlab-snu/PPI-Landscape-2025. The codes for ProteinMPNN-based productivity rescue strategy are available on GitHub at https://github.com/CSSB-SNU/ab-expression-rescue.

## Acknowledgements

This research was supported by the Bio&Medical Technology Development Program of the National Research Foundation (NRF) funded by the Korean government (MSIT) (to T.-Y.Y., grant number RS-2024-00397865).

## Author contributions

T.-Y.Y. and M.B. conceived of and supervised the project. C.C., B.-K.S., J.H.J., and H.K. designed the experiments. C.C., B.-K.S., J.H.J., J.P., J.L., S.C., J.C.^1^, H.C., B.Y., S.B.L., and C.Y.L. performed experiments using the SPID platform. H.K. and M.B. developed the AI-guided design strategies and performed the associated computational modeling. H.-T.A., J.E.K., Y.B., and Y.-Y.C. generated animal models and performed animal experiments. C.C., B.-K.S., J.H.J., H.K., H.-T.A., J.P., B.C., H.L., J.C.^2^, M.B. and T.-Y.Y. analyzed all data. C.C., B.-K.S., H.K., H.-T.A., and J.P. visualized all data. C.C., B.-K.S., H.K., M.B., and T.-Y.Y. wrote the manuscript with input from all other authors.

## Extended Data Figures

**Extended Data Fig. 1.**
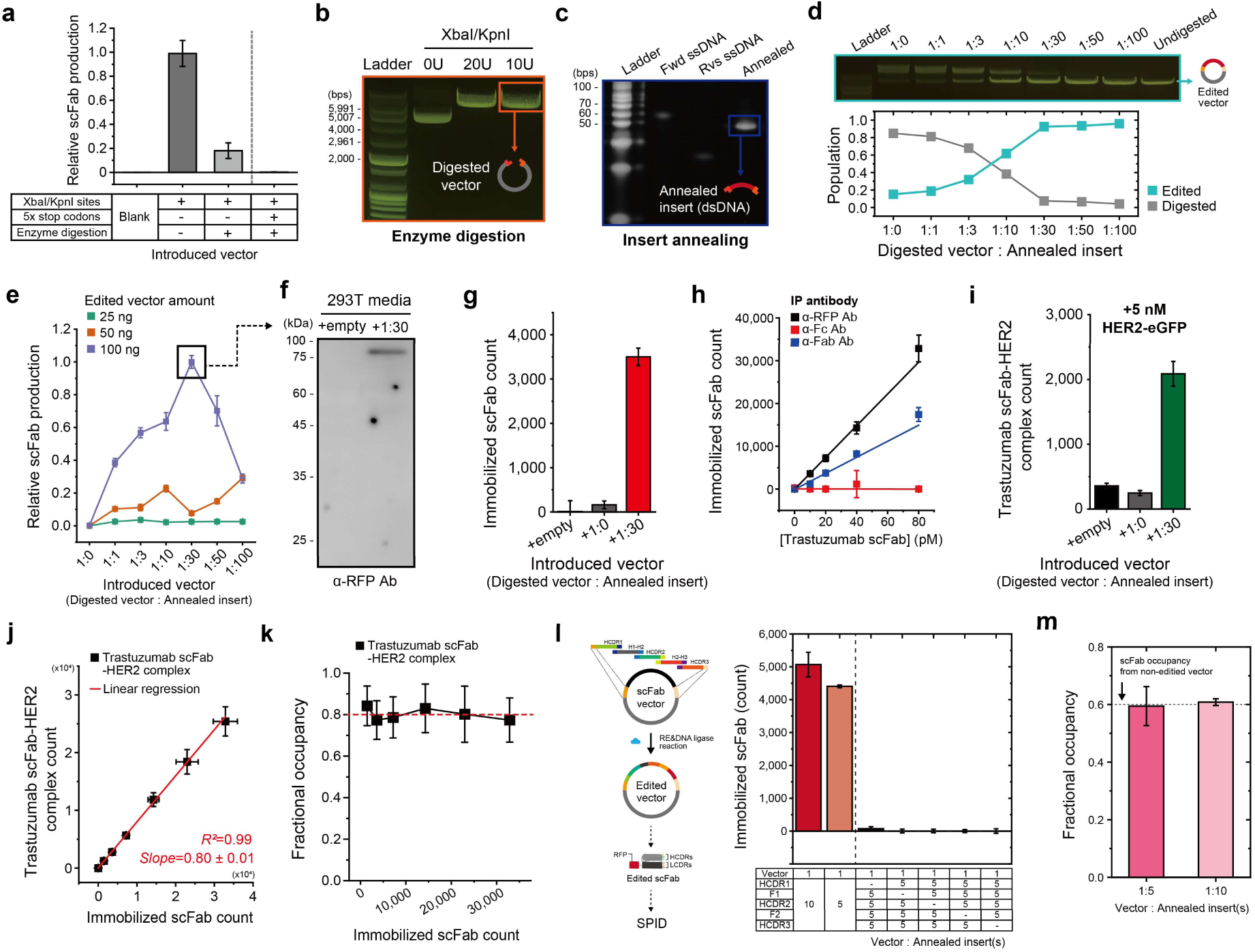
Optimization and validation of the high-throughput modular CDR editing workflow for SPID analysis. **a**, Complete inhibition of scFab production via the introduction of a five-tandem stop codon cassette within the HCDR3. **b**, Restriction enzyme digestion efficiency of the modular trastuzumab scFab vector. Simultaneous digestion with *KpnI* and *XbaI* achieves >99% purity of digested vectors. **c,** Annealing efficiency between forward and reverse ssDNAs, showing >99% purity of the resulting dsDNA product. **d,** *In vitro* ligation efficiency measured across various ratios of annealed inserts to digested vector. **e**, Expression levels of trastuzumab scFabs in HEK293T cells after two days of expression according to the introduced ligated plasmid. **f,** Validation of secreted trastuzumab scFabs generated via modular CDR editing. **g**, Detection and single-molecule quantification of secreted trastuzumab scFabs using SPID. **h**, Single-molecule counting of surface-immobilized trastuzumab scFabs within the FOV using the indicated IP antibodies. **i**, Specific binding between secreted trastuzumab scFabs and HER2-eGFP. 5 nM of HER2-eGFP was loaded onto the surface-immobilized scFab as shown in (**g**). **j,** Linear correlation between the trastuzumab scFab-HER2 complex count and the density of surface-immobilized trastuzumab scFabs. **k,** Consistent fractional occupancies across increasing concentrations of surface-immobilized trastuzumab scFabs. **l**, Schematic overview and production of pembrolizumab scFab via SPID-based multiple CDR editing. **m,** Specific binding between secreted pembrolizumab scFabs and PD-1-eGFP. 5 nM of PD-1-eGFP was loaded onto surface-immobilized scFab as shown in (**l**).

**Extended Data Fig. 2.**
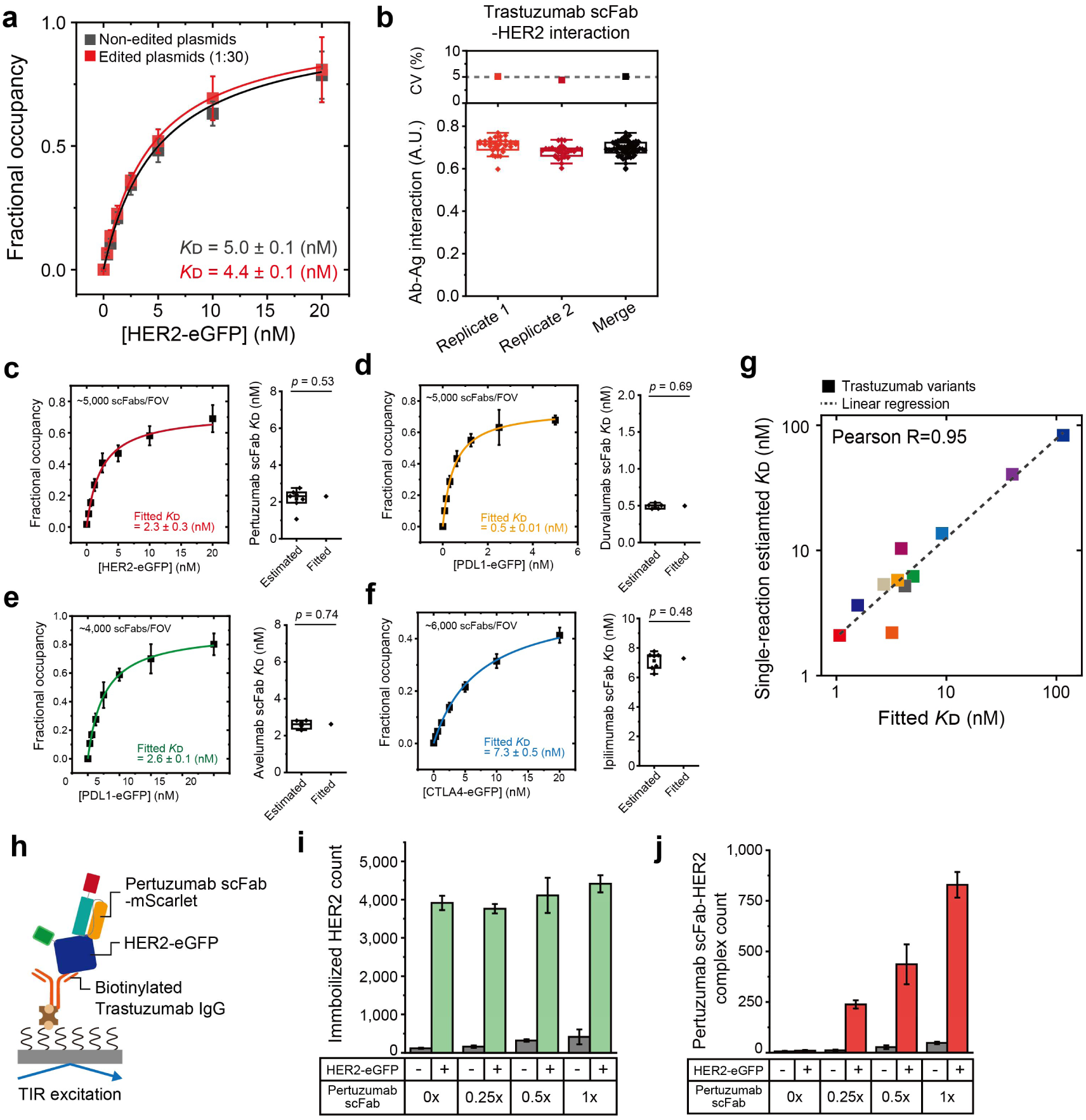
Validation of SPID robustness and versatility across diverse antibody-antigen pairs. **a,** Comparison of the binding curves of trastuzumab scFab-HER2 interactions generated via different scFab production methods. **b**, Repeatability of trastuzumab scFab production and binding affinity measurement. Thirty independent CDR editing were performed to reconstitute the original HCDR3 sequence in each replicate trial (sixty in total). Intra-chip coefficients of variation (CVs) are displayed. **c-f,** Binding curves for eGFP-labeled antigens to their corresponding scFabs used for *K*_D_ fitting. *K*_D_ values obtained by single-reaction estimates and curve fitting are compared (Two-sided one-sample t-test). (**c**) Pertuzumab scFab-HER2, (**d**) Durvalumab scFab-PD-L1, (**e**) Avelumab scFab-PD-L1, (**f**) Ipilimumab scFab-CTLA-4. **g**, Correlation between single-reaction estimated *K*_D_ values (modular CDR editing) and fitted *K*_D_s (conventional cloning) across 11 different trastuzumab scFab variants. **h,** Schematic of co-immunoprecipitation of pertuzumab scFab and trastuzumab IgG. **i,** Surface immobilization of HER2-eGFP by biotinylated trastuzumab IgG using SPID, demonstrating independence from pertuzumab scFab introduction. **j,** Simultaneous binding of pertuzumab scFab and trastuzumab IgG to HER2-eGFP.

**Extended Data Fig. 3.**
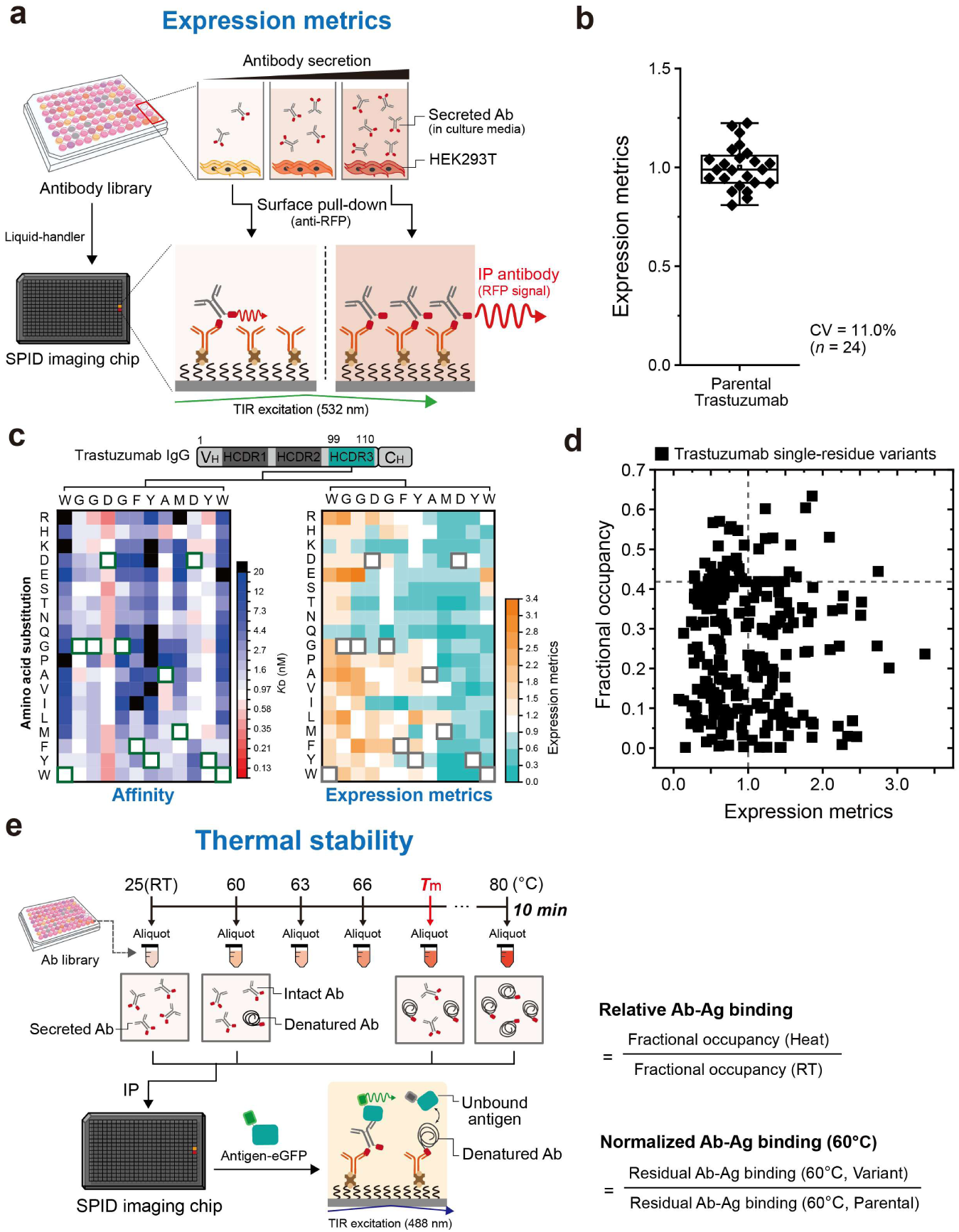
Multi-parametric characterization of antibody developability via SPID. **a,** Schematic for measuring expression metrics using SPID. **b,** Repeatability of expression metrics. Twenty-four independent CDR edits were performed to reconstitute trastuzumab scFab in each trial. Intra-chip CV is displayed. **c**, DMS heatmaps of trastuzumab IgG variants, illustrating the landscapes of antigen binding affinity (*K*_D_) and expression metrics. **d,** Correlation between fractional occupancies and expression metrics for trastuzumab IgG single-residue variants (*n*=380). **e,** Schematic for assessing antibody thermal stability based on residual antigen-binding capacity after heat treatment.

**Extended Data Fig. 4.**
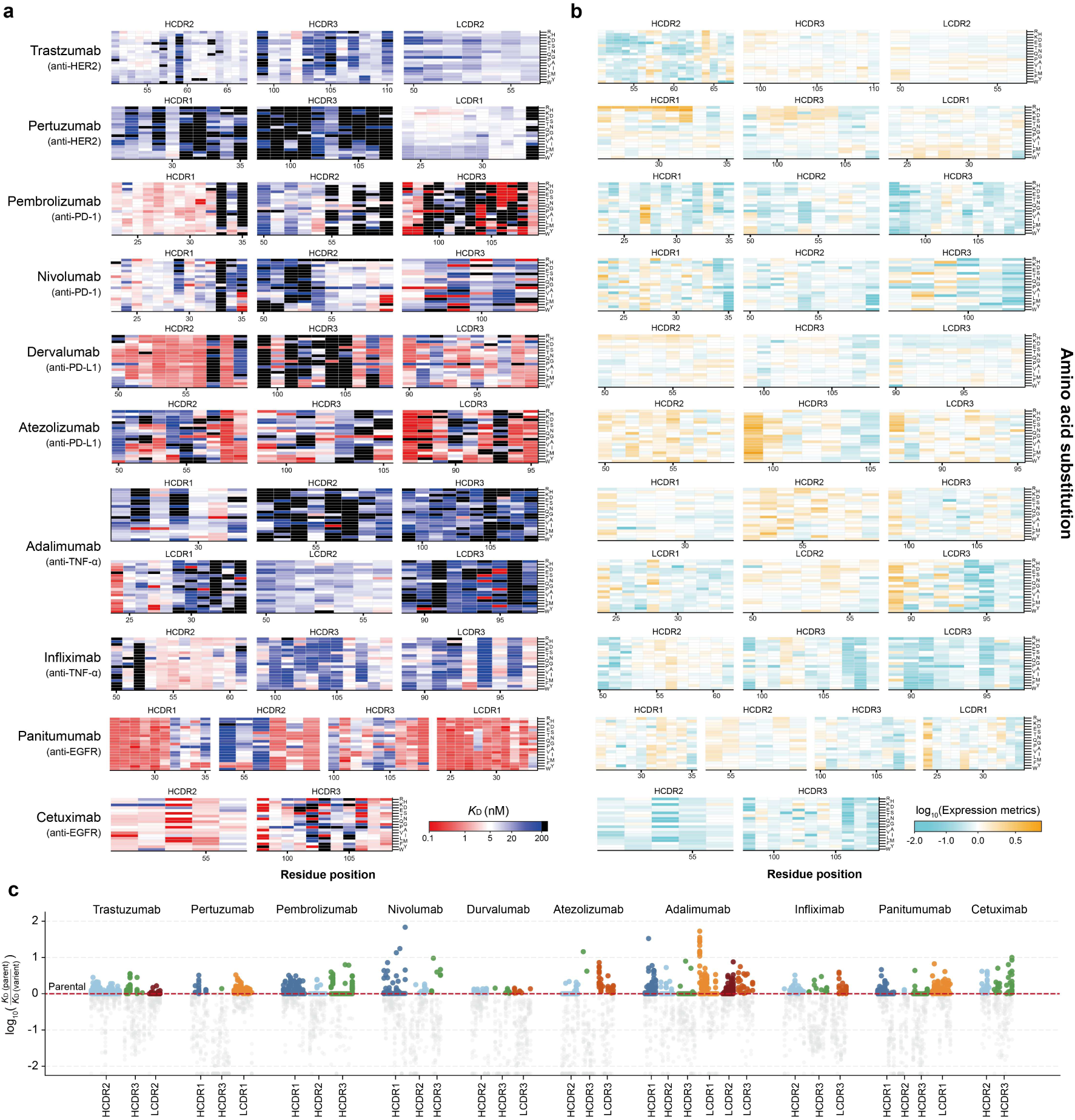
Comprehensive DMS landscapes of binding affinity and expression for ten therapeutic antibodies. **a,b,** High-throughput DMS heatmaps for ten FDA-approved therapeutic antibodies, showing fold changes in (**a**) antigen-binding affinity (*K*_D_) and (**b**) expression metrics. All data are normalized to their respective parental sequences. **c,** CDR-specific affinity distributions. Only affinity-permissive or enhancing variations relative to the parental baseline are shown in color, while deleterious mutations are indicated in gray.

**Extended Data Fig. 5.**
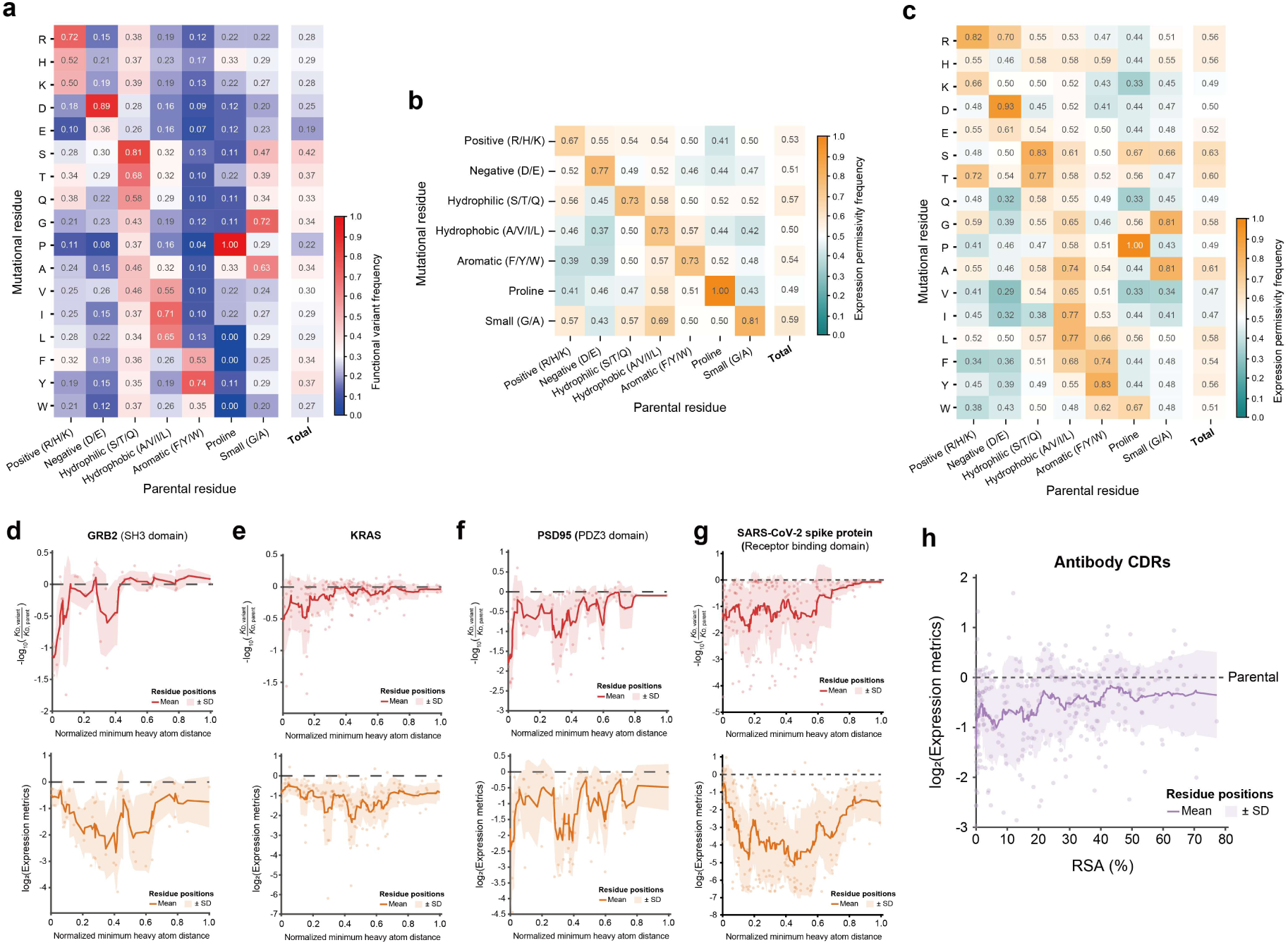
Physicochemical and structural determinants of mutational permissivity. **a**, Functional variant frequency categorized by amino acid type. **b,c,** Expression permissivity frequency categorized by (**b**) physicochemical property and (**c**) amino acid type. **d-f,** Distance-dependent functional profiles for individual globular proteins. Mean functional changes are plotted as a function of the minimum heavy atom distance from the binding interface for (**d**) SH3 domain of GRB2, (**e**) KRAS, (**f**) PDZ3 domain of PSD95, and (**g**) RBD of SARS-CoV-2 Spike protein. **h,** Relationship between of RSA and expression metrics for 324 antibody CDR residues.

**Extended Data Fig. 6.**
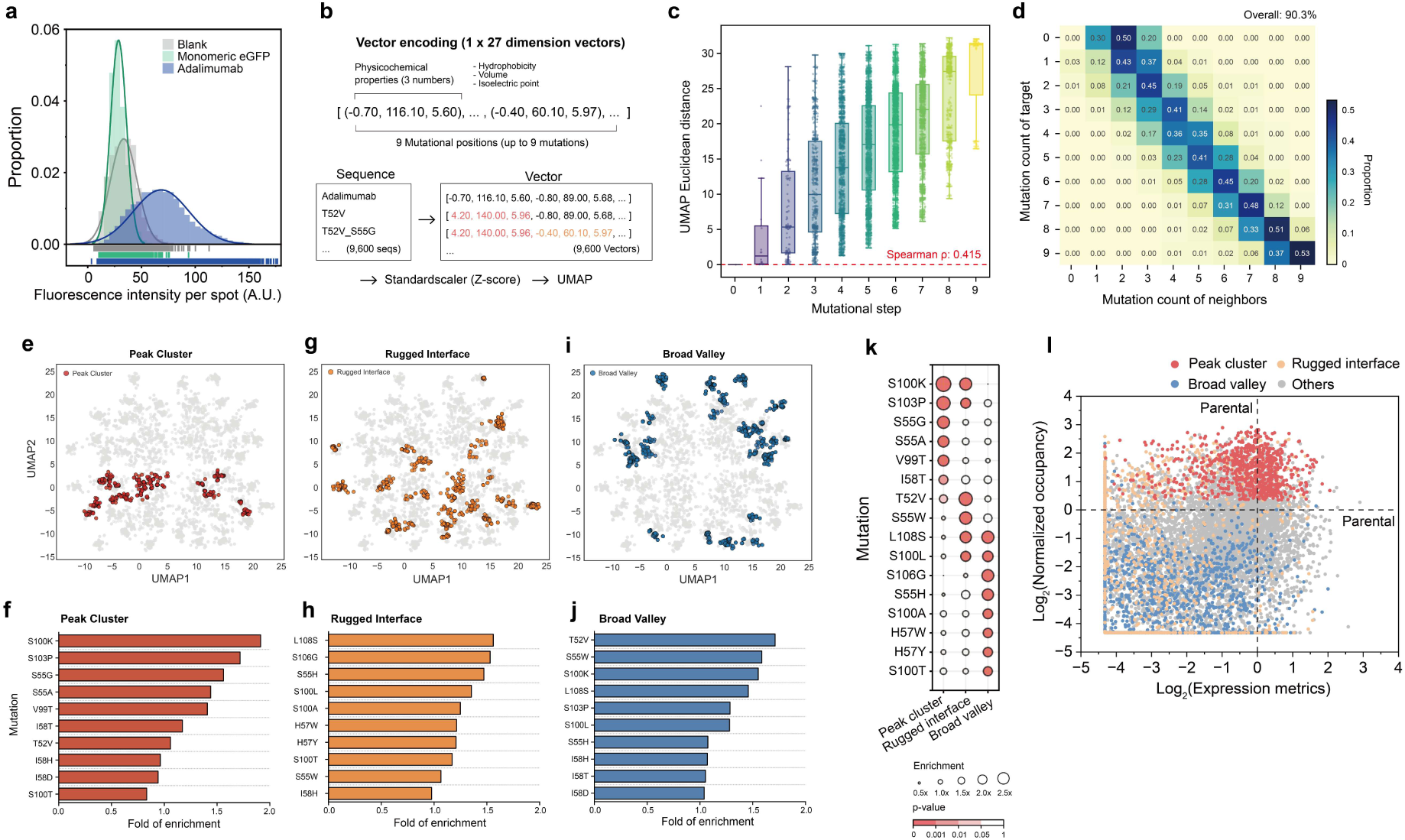
Physicochemical embedding and topographical characterization of the sequence space. **a,** Fluorescence distribution of single adalimumab–target complexes. Rightward fluorescence shifts per spot indicate preferential binding of adalimumab to trimeric TNF-α over the monomeric form (eGFP). **b,** Schematic of sequence vectorization, where each variant is encoded as a 27-dimensional physicochemical vector (hydrophobicity, volume, isoelectric point) before UMAP projection. **c,** Correlation between mutational steps and UMAP Euclidean distance, showing spatial expansion with increasing mutational load (Spearman’s ρ = 0.415). **d,** Heatmap correlating mutation counts between each sequence and its UMAP neighbors, demonstrating high local topographical consistency (90.3%). **e–j,** Topographical distribution and mutational enrichment across three major clusters: **(e** and **f)** peak cluster, **(g** and **h)** rugged interface, and **(i** and **j)** broad valley. Histograms show fold enrichment of single mutations within each cluster. **k,** Bubble chart summarizing mutational enrichment across topographical groups, with bubble size indicating enrichment magnitude and color denoting significance (p-value). **l,** Fitness scatter plot (expression metric versus relative occupancy) color-coded by topographical assignment: peak cluster (red), rugged interface (brown), broad valley (blue), and others (gray).

**Extended Data Fig. 7.**
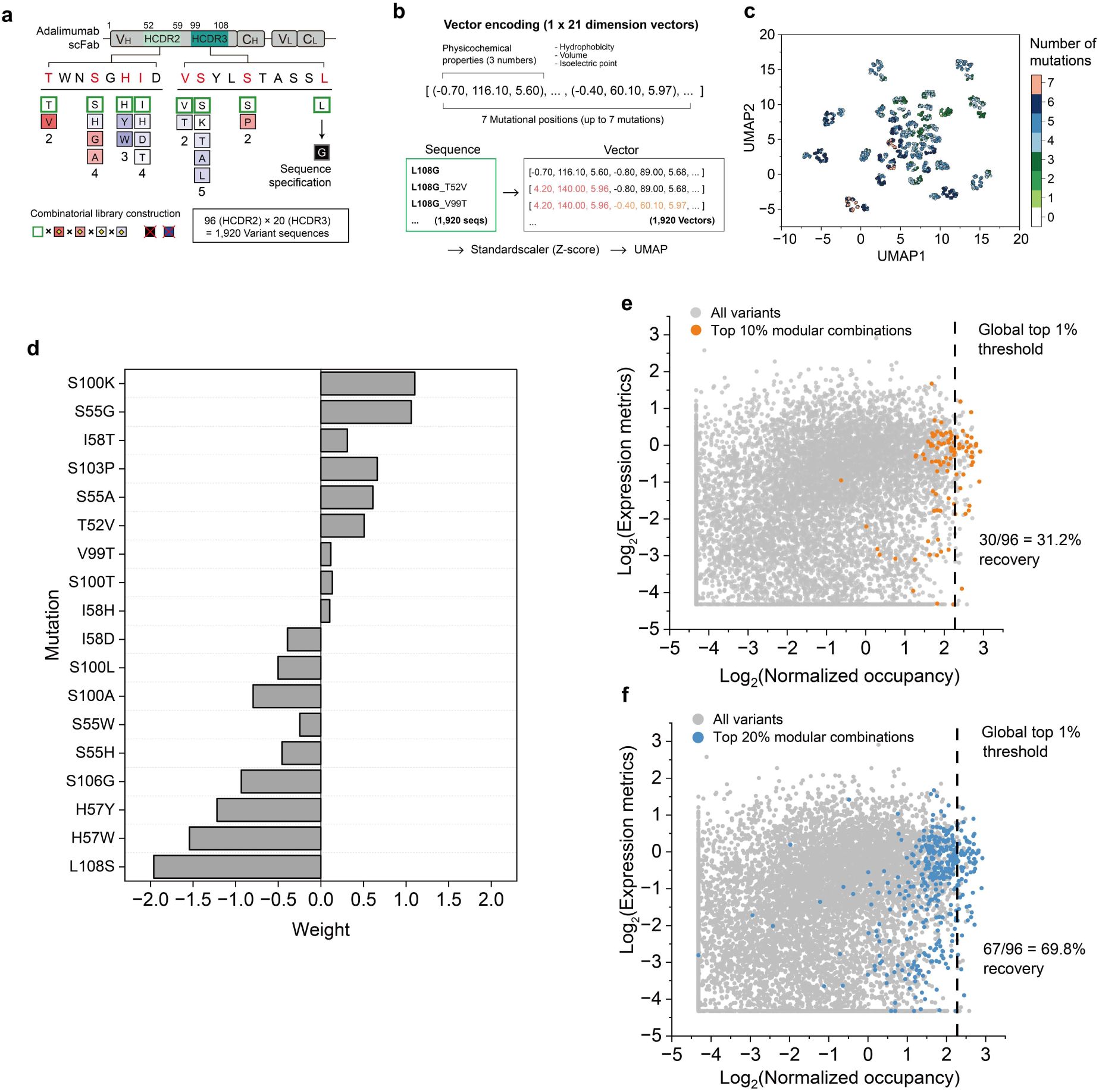
Design, physicochemical embedding, and fitness evaluation of the L108G combinatorial library. **a,** Schematic overview of library construction on the L108G fixed background. Substitutions across seven targeted positions in HCDR2 and HCDR3 generated 1,920 variants. **b,** Schematic of sequence vectorization, where each variant was encoded as a 21-dimensional continuous vector reflecting hydrophobicity, volume, and isoelectric point across the seven mutated sites, followed by Z-score normalization and UMAP projection. **c,** Two-dimensional UMAP plot of the 1,920 variants, with points color-coded by mutation count. **d,** Additive weights of individual single-amino-acid mutations derived from a ridge regression model. **e,f,** Fitness scatter plots (expression metrics versus normalized occupancy) illustrating the modular assembly strategy. Variants recovered by combinatorially pairing the top (**e**) 10% (orange) or (**f**) 20% (blue) modules selected independently from each CDR are highlighted. Dashed lines mark the global top 1% fitness threshold, with recovery rates of 31.2% and 69.8%, respectively.

**Extended Data Fig. 8.**
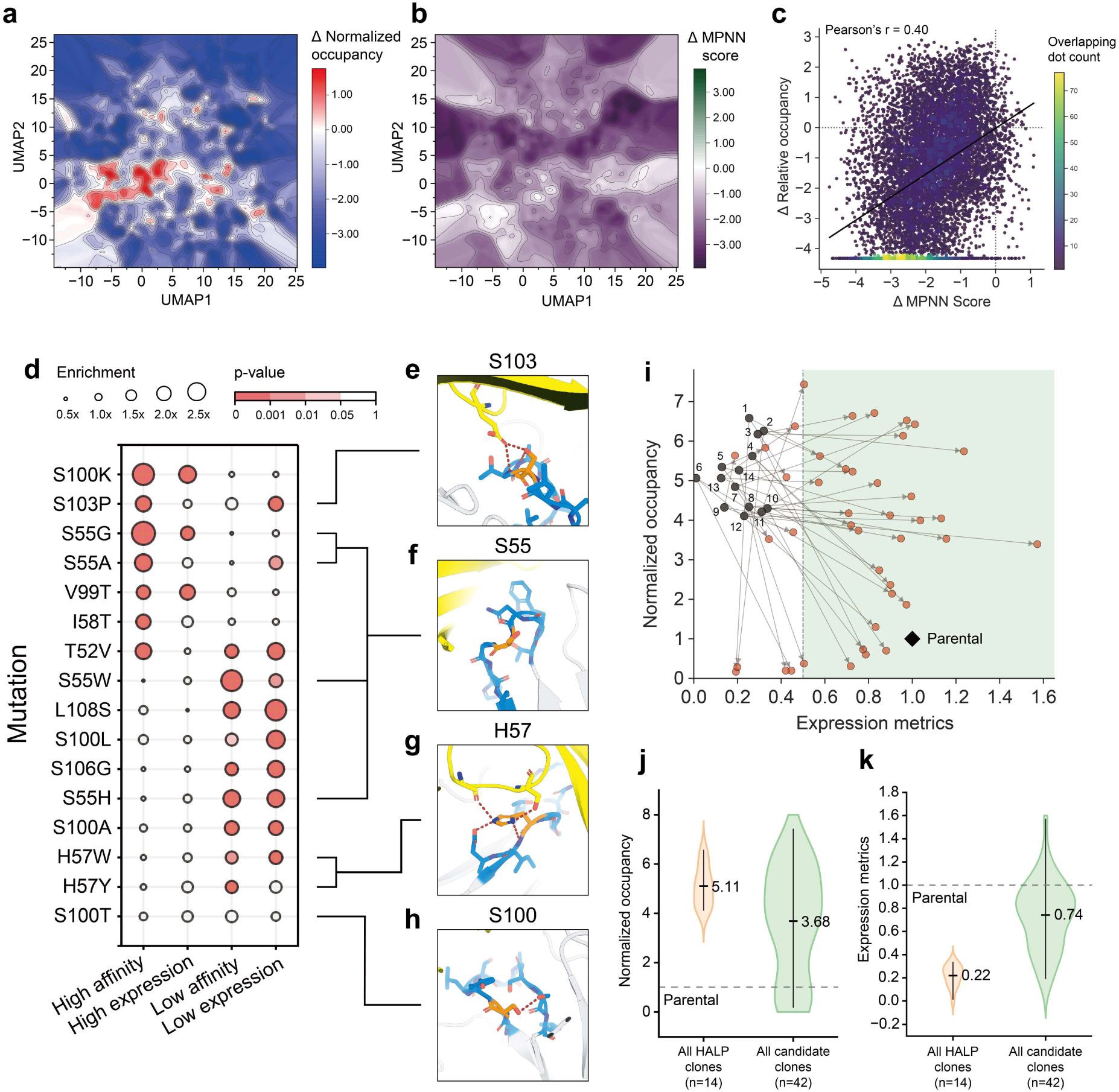
Topographical affinity-stability relationships and broad validation of the computational rescue strategy. **a,b,** 2D contour landscapes illustrating the deviation (Delta) of (**a**) the experimental relative occupancy and (**b**) the ProteinMPNN compatibility score relative to the parental clone. Color gradients indicate variations from the parental baseline (white). **c,** Correlation scatter plot between the relative occupancy and MPNN score (Pearson’s r = 0.40). Data point density is visualized using a count-based color scale to reflect overlapping dot counts. **d,** Bubble chart quantifying the enrichment and statistical significance (p-value) of specific single mutations across distinct fitness quadrants (high affinity/high expression versus low affinity/low expression). Bubble size denotes the fold of enrichment. **e–h,** Structural snapshots of the parental complex highlighting the local interaction environments of key residues associated with low expression fitness profiles: (**e**) S103 in HCDR3 at the antigen-binding interface, forming hydrogen bonds with both the antigen and neighboring HCDR3 residues, (**f**) S55 in HCDR2 and its local structural environment, (**g**) H57 in HCDR2, forming hydrogen bonds with neighboring HCDR2 residues while simultaneously contacting the antigen, and (**h**) S100 in HCDR3, forming intra-loop polar contacts with neighboring CDR3 residues. In all panels, the antigen is shown in yellow, the antibody framework in white, HCDR2 and HCDR3 in blue, and the target residue in orange. Key interactions are indicated by brick-colored dashed lines. **i,** Two-dimensional fitness trajectory plot tracking the comprehensive rescue outcomes for all 14 HALP clones and 42 candidate clones. Directional arrows connect the initial HALP clones (numbered gray dots) to their corresponding candidates (red dots). **j,k,** Violin plots comparing the distributions and mean values of (**j**) relative occupancy and (**k**) the expression metric between the initial HALP clones (n=14) and the generated pool of candidate clones(n=42). Dashed lines indicate the parental baseline levels.

**Extended Data Fig. 9.**
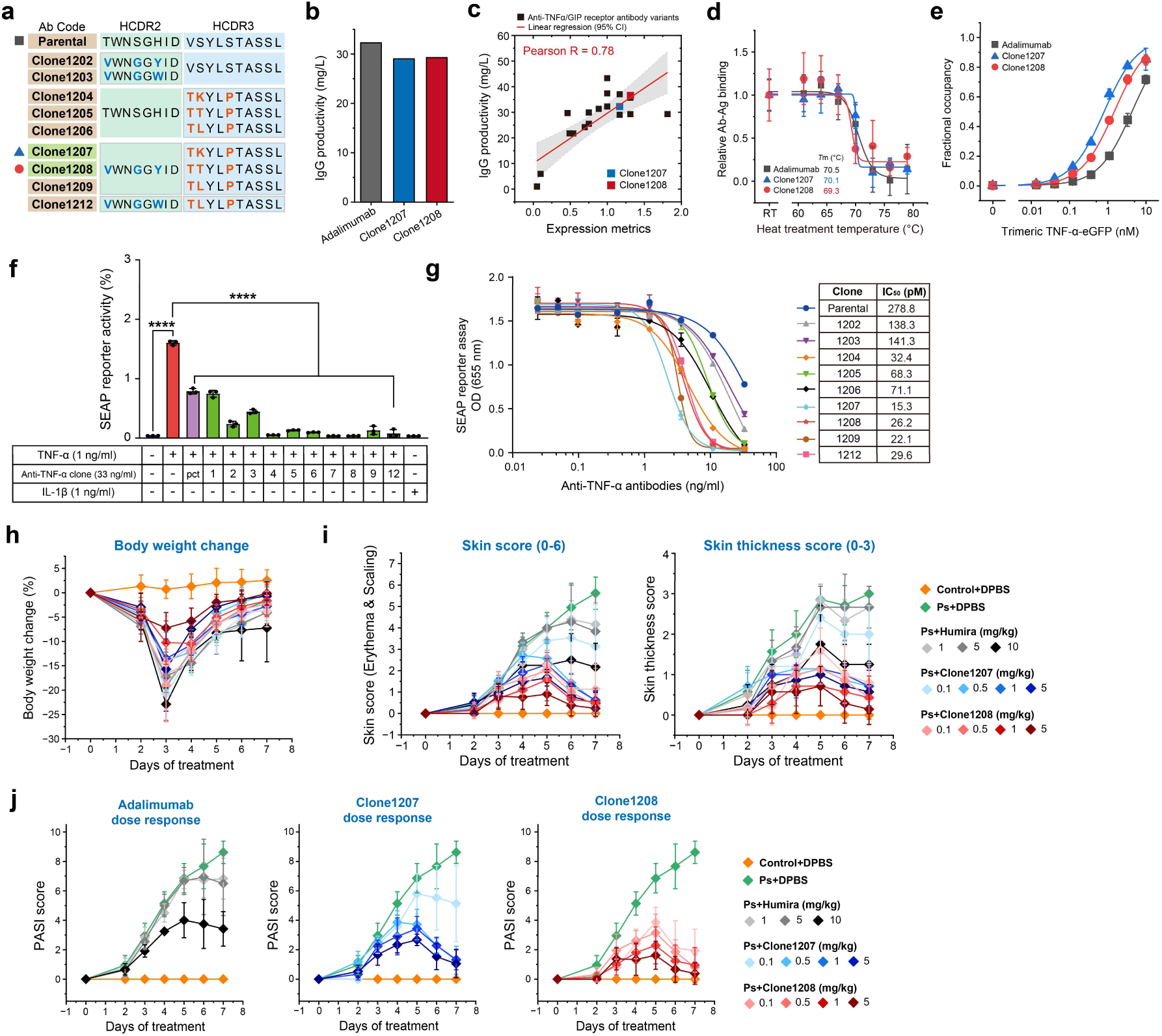
Molecular characterization and *in vivo* validation of engineered adalimumab variants. **a**, HCDR2 and HCDR3 sequences of the parental adalimumab and the engineered variants. **b-e,** Comprehensive profiling of engineered adalimumab variants. (**b**) Volumetric productivity of parental adalimumab and clones 1207 and 1208, purified from 40 ml HEK293F cultures. (**c**) Correlation between SPID-measured expression metrics and productivity across large-scale (up to 10 liters) cultures of anti-TNF-α or anti-GIPR antibodies in HEK293F cells. Data points for clones 1207 and 1208, produced in 7L cultures, are explicitly highlighted. (**d**) Thermal stability assessment via melting curve analysis. (**e**) Dose-dependent binding curves with trimeric TNF-α with SPID. **f,** *In vitro* neutralization activity assessed using an NF-κB–responsive SEAP reporter assay. **g,** Concentration-response analysis and calculated *IC*₅₀ values demonstrate the enhanced neutralizing potency of the engineered clones. **h,** Body weight change (%) of mice during the experiment, indicating no significant treatment-related toxicity. **i,** Detailed PASI score components (skin score (erythema and scaling) and thickness) plotted over the 7-day treatment period. **j,** Dose-response relationship for adalimumab, clone1207, and clone1208 based on PASI scores at day 7.

**Extended Data Fig. 10.**
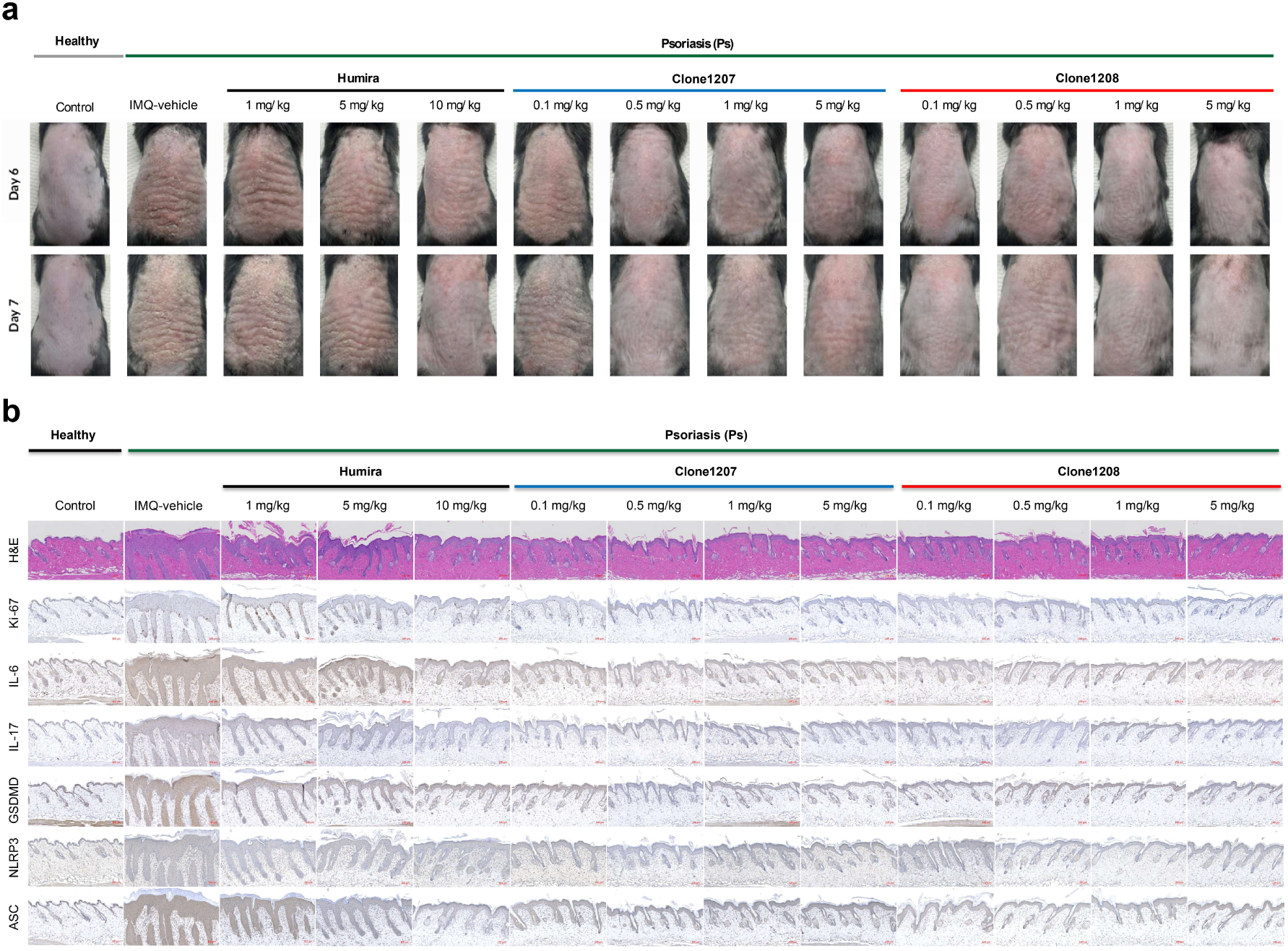
Comprehensive macroscopic and microscopic analysis of skin tissue from the psoriasis mouse model. **a,** Representative photographs of dorsal skin on day 6 and day 7 from the imiquimod (IMQ)-induced psoriasis model, showing the dose-dependent therapeutic effects of each antibody. **b,** Representative images of histological and immunohistochemical staining of dorsal skin sections collected on day 7 from the IMQ-induced psoriasis model. The panels show the dose-dependent effects of adalimumab, clone1207, and clone1208 on key pathological features of psoriasis. Staining includes H&E for epidermal hyperplasia, Ki-67 for keratinocyte proliferation, IL-6 and IL-17 for key inflammatory cytokines, and GSDMD, NLRP3, and ASC as markers for the pyroptosis and inflammasome pathways. The results demonstrate that the engineered variants, particularly clone1208, effectively suppress these markers of inflammation and hyperproliferation.

