## Supplementary Information for "Landscape-scale navigation unlocks antibody CDR structural logic for AI-guided rescue and therapeutic optimization"

**List of contents:**
Supplementary Notes 1-2

**Supplementary Notes**

**Supplementary Note 1 | Structural analysis of expression-sensitive positions**

To examine whether the expression sensitivity observed in the DMS data could be structurally rationalized, we analyzed the local structural environments of residues frequently mutated in the bottom 10% of expression levels (Fig. 4d). If sequence-structure compatibility contributes to CDR expression, mutations at these positions should be explainable by disruption of specific structural interactions. We focused on eight representative residues across HCDR2 and HCDR3, four of which are presented in the main figures (T52, I58, S106, L108; Fig. 4e-h) and four in the extended data figures (H57, S55, S100, S103; Extended Data Fig. 8).

T52 in HCDR2 forms an intra-loop hydrogen bond network through its hydroxyl group with neighboring HCDR2 residues, maintaining the loop conformation proximal to the antigen-binding interface (Fig. 4e). The expression loss observed for T52V is consistent with disruption of this polar network, suggesting that hydrogen bonding capacity at this position contributes to proper loop scaffolding.

I58 anchors HCDR2 to the framework through hydrophobic packing against the framework β-sheet surface (Fig. 4f). The strong destabilizing effects of I58D and I58H are consistent with the introduction of polar or charged residues into this hydrophobic interface. Notably, I58T is well-tolerated despite introducing a hydroxyl group, which can be explained by threonine's methyl group partially satisfying the hydrophobic interface when oriented toward the framework packing cavity. This suggests that expression is maintained as long as the substituted residue can present a hydrophobic surface toward the framework, regardless of whether it is strictly nonpolar.

S106 in HCDR3 participates in a dense intra-loop hydrogen bond network that scaffolds the loop conformation without directly contacting the antigen (Fig. 4g). The destabilizing effect of S106G is consistent with simultaneous loss of hydroxyl-mediated hydrogen bonds and introduction of backbone flexibility at this position.

L108 at the C-terminal stem of HCDR3 packs hydrophobically against the antibody framework, mirroring the structural role of I58 in HCDR2 (Fig. 4h). L108S is among the most expression-destabilizing mutations observed, consistent with the introduction of polarity into a hydrophobic cavity.

We further examined S103, S55, H57, and S100, whose structural environments are described in the Extended Data Fig. 8. S103 at the antigen-binding interface of HCDR3 contributes to both antigen contact and local loop conformation through side-chain hydrogen bonds with the antigen and neighboring residues (Extended Data Fig. 8e). The strong destabilizing effect of S103P is consistent with proline's rigid cyclic geometry imposing backbone dihedral angles incompatible with the native loop conformation, compounded by the loss of backbone NH hydrogen bonding capacity.

S55 in HCDR2 presents a less readily explained pattern (Extended Data Fig. 8f). While S55G is well-tolerated, S55H, S55W, and S55A are destabilizing without a clear physicochemical trend, suggesting a complex interplay between loop geometry, antigen contact, and steric constraints that cannot be attributed to a single dominant mechanism. This position may also be particularly susceptible to epistatic interactions with neighboring residues, making it difficult to isolate the structural effect of individual substitutions.

H57 in HCDR2 forms a hydrogen bond network with neighboring residues while simultaneously contacting the antigen, with its imidazole side chain positioned to fulfill both roles (Extended Data Fig. 8g). The strong destabilizing effect of H57W is consistent with the bulky indole disrupting the precise local geometry, whereas the partial tolerance of H57Y can be explained by tyrosine's ability to partially recapitulate the native polar interactions through its hydroxyl group.

S100 in HCDR3 stabilizes the loop conformation through intra-loop polar contacts with neighboring residues rather than direct antigen engagement (Extended Data Fig. 8h). The destabilizing effects of S100L and S100A, alongside the tolerance of S100K and S100T, indicate that hydrogen bonding capacity is the key requirement at this position.

Collectively, seven of the eight positions examined show mutational sensitivity that is directly explicable by one of two structural mechanisms: disruption of intra-loop hydrogen bond networks that scaffold loop geometry (T52, H57, S100, S103, S106), or introduction of polarity into hydrophobic CDR-framework interfaces (I58, L108). The single exception, S55, likely reflects more complex context-dependent effects. These observations provide structural support for the notion that sequence-structure compatibility contributes to CDR expression, consistent with the principles that underlie folding and expression in globular proteins.

**Supplementary Note 2 | Structural analysis of expression-rescued candidates**

To understand the structural basis of the expression improvements described in Fig. 4o-q, we examined the local structural environments of the rescue substitutions in HALP clones 2, 3, and 8, each of which achieved substantial expression recovery through a single amino acid change while maintaining high antigen affinity.

In HALP clone 2, the rescue substitution reverts S106G to serine, restoring the hydroxyl-mediated hydrogen bond network within the HCDR3 loop (Fig. 4o). This recovers the intra-loop polar scaffold that scaffolds HCDR3 loop geometry, improving expression to 1.24-fold while maintaining a 5.74-fold affinity gain. That restoration of this single network is sufficient to substantially rescue expression, even with I58D remaining, highlights the independent contribution of the HCDR3 intra-loop network to the overall expression phenotype.

In HALP clone 3, the rescue substitution I58D→I58T partially restores the hydrophobic HCDR2-framework interface. Although threonine is polar, its methyl group can satisfy the hydrophobic packing cavity when oriented toward the framework surface (see B1), recovering the CDR-framework contact disrupted by I58D. This single change improves expression to 0.73-fold to parental adalimumab while maintaining a 6.64-fold affinity gain (Fig. 4p), demonstrating that partial restoration of this interface is sufficient to meaningfully rescue productivity.

In HALP clone 8, the rescue substitution V99T introduces a hydroxyl group that forms a new hydrogen bond bridging the HCDR3 loop backbone to the antibody framework, stabilizing a CDR-framework junction that was insufficiently anchored in the source clone. This likely reduces the conformational entropy of the HCDR3 loop and suppresses misfolding propensity, recovering near-parental expression (1.00-fold) while maintaining a 4.60-fold affinity gain (Fig. 4q).

Collectively, these cases demonstrate that the expression defects of affinity-matured clones can be selectively corrected by single substitutions that either restore disrupted structural interactions or introduce new stabilizing contacts at CDR-framework interfaces, without sacrificing the affinity gains that required extensive combinatorial optimization to achieve.
